# Limited evidence for parallel evolution among desert adapted *Peromyscus* deer mice

**DOI:** 10.1101/2020.06.29.178392

**Authors:** Jocelyn P. Colella, Anna Tigano, Olga Dudchenko, Arina D. Omer, Ruqayya Khan, Ivan D. Bochkov, Erez L. Aiden, Matthew D. MacManes

**Affiliations:** University of New Hampshire, Molecular, Cellular, and Biomedical Sciences Department, Durham, NH 03824, USA; (JPC), (AT) (MDM); Hubbard Genome Center, University of New Hampshire, Durham, NH 03824, USA; Department of Computer Science, Department of Computational and Applied Mathematics; Center for Theoretical and Biological Physics, Rice University, Houston, TX 77030, USA; (OD), (ELA); The Center for Genome Architecture, Department of Molecular and Human Genetics, Baylor College of Medicine, Houston, TX 77030, USA; (ADO), (RK), (IDB); Department of Computer Science, Department of Computational and Applied Mathematics, Rice University, Houston, TX 77030, USA; Shanghai Institute for Advanced Immunochemical Studies, ShanghaiTech University, Shanghai 201210, China; School of Agriculture and Environment, University of Western Australia, Perth, WA 6009, Australia

**Keywords:** dehydration, desert, parallel evolution, *Peromyscus*, thermoregulation

## Abstract

Phenotypic plasticity enables an immediate response to changing conditions, but for most species, evolutionary change through adaptation will be more important for long-term survival. Warming climate and increasing desertification urges the identification of genes involved in heat-and dehydration-tolerance to better inform and target biodiversity conservation efforts. Comparisons among extant desert adapted species can highlight parallel or convergent patterns of genome evolution through the identification of shared signatures of selection. We generate chromosome-level genome assembly for the canyon mouse (*Peromyscus crinitus*) and test for signature of parallel evolution by comparing signatures of selective sweeps across population-level genomic resequencing data from another desert specialist deer mouse (*P. eremicus*) and a widely-distributed habitat generalist (*P. maniculatus*), that may locally adapted to arid conditions. We identify few shared candidate loci involved in desert adaptation and do not find support for a shared pattern of parallel evolution. Instead, we hypothesize divergent molecular mechanisms of desert adaptation among deer mice, potentially tied to species-specific historical demography, which may limit or enhance adaptation. We identify a number of candidate loci experiencing selective sweeps in the *P. crinitus* genome that are implicated in osmoregulation (*Trypsin, Prostasin*) and metabolic regulation (*Kallikrein, eIF2-alpha kinase GCN2, APPL1/2*), which may be important to accommodating hot and dry environmental conditions.

## INTRODUCTION

Increasing global temperatures and altered patterns of precipitation threaten biodiversity worldwide (Moritz et al. 2008; Cahill et al. 2013; Urban 2015). Phenotypic plasticity enables an immediate response to changing conditions but adaptation through evolutionary change will be critical for the long-term survival of most species (Hoffman and Sgrò 2011; Cahill et al. 2013).Range shifts upward in elevation and latitude have been documented in a number of terrestrial species and interpreted as a response to warming (Chen et al. 2011; Tingley and Beissinger 2013; Freeman et al. 2018); however, responses vary even among closely-related species or populations (Hoffman and Willi 2008; Moritz et al. 2008). Geographic range shifts are often governed by the physiological limits of species, which are in part controlled by genetics and have been shaped by neutral and selective evolutionary forces across many generations. Population genomics methods enable genome-wide scans for selection to identify genes and molecular pathways that may be involved in local adaptation (Bassham et al. 2018; Garcia-Elfring et al. 2019). For species adapted to similar environments, parallel or convergent evolution can be inferred if a greater number of genes or phenotypes share signatures of selection than would be expected under a purely stochastic model of evolution (e.g., drift). The same gene or suite of genes consistently tied to a specific adaptive phenotype across distantly related taxa is consistent with a signal of convergent evolution. In contrast, for taxa that share a recent common ancestor, signatures of selection at the same loci may reflect parallel selection on either new mutations or shared ancestral variation or similar demographic histories, resulting in the same phenotypic effect. Evidence of convergent or parallel evolution can highlight common loci involved in shared adaptive phenotypes (Rundle et al. 2000; McDonald et al. 2009), while a lack of concerted evolution may identify idiosyncratic evolutionary strategies to achieve the same phenotypic result.

As a model taxon (Dewey and Dawson 2001; Bedford and Hoekstra 2015) inhabiting varied environments throughout North America, deer mice (genus *Peromyscus*) are a frequent and productive subject of adaptation studies (e.g., physiological, Storz 2007; behavioral, Hu and Hoekstra 2017; genetic, Cheviron et al. 2012; Storz and Cheviron 2016; Tigano et al. 2020). Physiological similarity of deer mice to lab mice (Mus musculus) further broadens the implications of evolutionary and ecological investigations of Peromyscus by linking to biomedical sciences. The genus Peromyscus (N = 67 species; mammaldiversity.org) is hypothesized to be the product of a rapid ecological radiation across North America (origin ∼8 Mya, radiation ∼5.71 Mya; Platt et al. 2015), evident in their varied ecological niches and rich species diversity (Glazier 1980; Riddle et al. 2000; Bradley et al. 2007; Platt et al. 2015; Lindsey 2020). Peromyscus display tremendous thermoregulatory plasticity and can be found in extreme thermal environments, ranging from cold, high elevations (Pierce and Vogt 1993; Cheviron et al. 2012, 2014; Kaseloo et al. 2014; Garcia-Elfring et al. 2019) to arid, hot deserts (Riddle et al. 2000; MacManes 2017; Tigano et al. 2020). Thermoregulation and dehydration tolerance are complex physiological traits, suggesting that several potential evolutionary routes could lead to the same phenotypic outcome. Within this framework, comparisons among divergent Peromyscus species adapted to similar environments may highlight shared adaptive polymorphisms or disparate evolutionary paths central to achieving the same phenotype (Cheviron et al. 2012; Ivy and Scott 2017; Hu and Hoekstra 2017; Storz et al. 2019). In cold environments, endotherms rely on aerobic thermogenesis to maintain constant internal body temperatures. Changes in both gene expression and the functional properties of proteins in deer mice (P. maniculatus) adapted to high-altitude suggest that changes in multiple hierarchical molecular pathways may be common in the evolution of complex physiological traits, such as thermoregulation (Wichman and Lynch 1991; Storz 2007; Cheviron et al. 2012; Storz and Cheviron 2016; Garcia-Elfring et al. 2019). Nonetheless, research focused on thermoregulatory adaptations in high-elevation species may be confounded by concurrent selection on other traits conferring fitness benefits, such as high hemoglobin oxygen-binding affinity (Storz and Kelly 2008; Storz et al. 2010; Natarajan et al. 2015), which is critically important given the low partial pressure of oxygen (PaO_2_) associated with high elevation environments. In hot environments, endotherms are challenged with balancing heat dissipation, energy expenditure, and water retention (Anderson and Jetz 2005), resulting in a different suite of behavioral, physiological, and molecular adaptations that enable survival (Schwimmer and Haim 2009; Degen 2012; Kordonowy et al. 2016), but may be confounded by acute or chronic dehydration. Understanding the biochemical mechanisms that enable survival under extreme environmental stress can provide important insights into the nature of physiological adaptation. Rapid environmental and ecological differentiation among *Peromyscus* species positions these small rodents as models for generating hypotheses surrounding species responses to accelerated warming (Cahill et al. 2013) and the potential for repeated adaptation to similar environments among closely related species. Numerous *Peromyscus* species are adapted to life in hot deserts, with each species and population subject to distinct histories of demographic variation and gene flow. These idiosyncratic histories have a direct impact on evolution, as effective population sizes (*N*_*e*_) are inextricably linked to the efficacy of selection and maintenance of genetic diversity in wild populations (Charlesworth 2009). Contemporary or historical gene flow may further help or hinder adaptive evolution through homogenization or adaptive introgression, respectively (Coyne and Orr 2004; Morjan and Reiseberg 2004; Jones et al. 2018; Tigano and Friesen 2016). Native to the American West, the canyon mouse (*P. crinitus*, Fig. 1) is a xerocole, highly specialized to life in hot deserts. In the lab, *P. crinitus* can survive in the absence of exogenous water, with urine concentration levels similar to that of desert-adapted kangaroo rats (*Dipodomys merriami*; Abbott 1971; MacMillen 1972; MacMillen and Christopher 1975; MacMillien 1983), but without equivalently specialized renal anatomy (Issaian et al. 2012). Canyon mice also exhibit a lower-than-expected body temperature relative to their size and can enter environmentally-induced torpor in response to drought, food limitation, or extreme external temperatures (McNab 1968; McNab and Morrison 1963; Morhardt and Hudson 1966; Johnson and Armstrong 1987), which facilitates survival in highly-variable and extreme desert environments. These phenotypes persist for multiple generations in the lab indicating they have a genetic basis (McNab and Morrison 1968). Cactus mice (*P. eremicus*) are frequently sympatric with *P. crinitus* and share the same adaptations described above for *P. crinitus* (Veal and Caire 1979; Kordonowy et al. 2017). Thus, we expect these two species to exhibit similar patterns of molecular adaptation. These two desert specialists belong to a monophyletic clade of deer mice, which also includes *P. merriami, P. californicus, P. eva*, and *P. fraterculus*, and is estimated to have diverged around 5-6 Mya (Platt et al. 2015). Other members of this clade exhibit similar adaptations to desert environments, including urine concentration, reduced water requirements, and environmentally-induced torpor (McNab and Morrison 1963; Veal and Caire 1979) suggesting that desert adaptation may represent the ancestral state of this clade. In contrast, the habitat generalist *P. maniculatus* (North American deer mouse) is phylogenetically basal to the two desert specialists examined here and has a geographically widespread distribution across North America. *Peromyscus maniculatus* inhabits a wide range of thermal environments, including hot southwestern deserts and cool, high elevations, but desert specialists are not its closest relatives and the species is not generally considered a xerocole. Locally adapted desert populations of *P. maniculatus* (subspecies *P. m. sonoriensis*), however, may exhibit patterns of selection similar to that of desert specialists, either through the parallel selective retention of functional ancestral polymorphisms or convergent selection on new mutations. Whole-genome assemblies are publicly available for both *P. eremicus* (PRJEB33593, ERZ119825; Tigano et al. 2020) and *P. maniculatus* (GCA_003704035.1), which positions these species as ideal comparatives against *P. crinitus* to identify genes and regulatory regions associated with desert adaptation, including those unique to desert specialists *P. eremicus* and *P. crinitus*.

**Figure 1.**
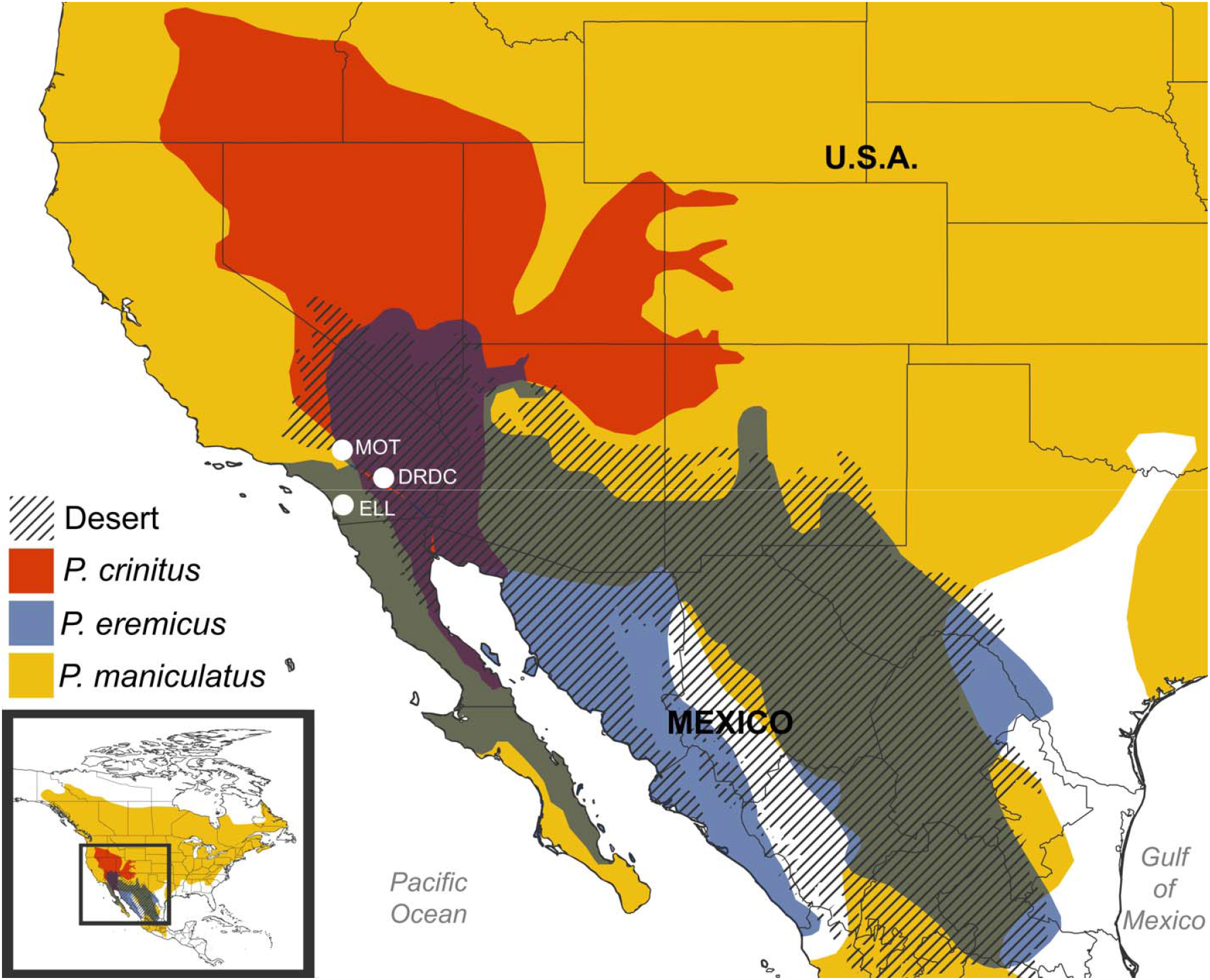
Geographic ranges of the three *Peromyscus* species examined in this study with major southwestern North American deserts denoted by diagonal hashing. *P. crinitus* range is in red, *P. eremicus* in blue, and *P. maniculatus* in yellow. Areas of sympatry denoted by color overlap: dark purple = yellow + red + blue and green = yellow + blue. MOT = Motte Rimrock Reserve, ELL = Elliot Chaparral reserve, DRDC = Philip Boyd Deep Canyon Desert Research Center.

Here, we investigate genomic signatures of selection among desert adapted *Peromyscus*. We contrast signatures of selective sweeps across three related *Peromyscus* species, two desert specialists (*P. crinitus* and *P. eremicus*) and one habitat generalist collected in an arid environment (*P. maniculatus*). We hypothesize that similar genes or functional groups will be under selection in related desert specialist species (*P. eremicus* and *P. crinitus*), due to their shared recent common ancestor and mutual association with hot, arid environments. In contrast, we hypothesize that *P. maniculatus* will show idiosyncratic evolutionary responses, with arid adaptation in this clade having evolved independently in response to local conditions. If similar genes and pathways are under selection in all three species, it would suggest local adaptation of *P. maniculatus* to desert conditions, and potentially, parallel or convergent evolution among divergent *Peromyscus* clades. Given the evolutionary distance of *P. maniculatus* to the two desert adapted species, a shared signature of selection across all three species may also indicate that adaptive responses to desert conditions are predictable and can occur repeatedly and potentially on short evolutionary timescales. Finally, we place selective sweep analyses into an evolutionary framework to interpret the varied evolutionary trajectories available to small mammals to respond to changing environmental conditions and to account for demographic and gene flow events.

## MATERIALS AND METHODS

### De novo genome sequencing and assembly

Wild mice were handled and sampled in accordance with the University of New Hampshire and University of California Berkeley’s Institutional Animal Care and Use Committee (130902 and R224-0310, respectively) and California Department of Fish and Wildlife (SC-008135) and the American Society of Mammalogists best practices (Sikes and Animal Care and Use Committee of the American Society of Mammalogists 2016).

For the assembly of the *P. crinitus* genome, DNA was extracted from a liver subsample from an individual collected in 2009 from the Philip L. Boyd Deep Canyon Desert Research Center (DRDC) in Apple Valley, California. To generate a high-quality, chromosome-length genome assembly for this individual we extracted high-molecular-weight genomic DNA using a Qiagen Genomic-tip kit (Qiagen, Inc., Hilden, Germany). A 10X Genomics linked-reads library was prepared according to the manufacturers protocol and sequenced to a depth of 70X on a HiSeq 4000 (Novogene, Sacramento, California, USA). 10X Genomics reads were *de novo* assembled into contigs using *Supernova* 2.1.1 (Weisenfeld et al. 2017). To arrange scaffolds into chromosomes, a Hi-C library for *P. crinitus* was constructed and sequenced from primary fibroblasts from the T.C. Hsu Cryo-Zoo at the University of Texas MD Anderson Cancer Center. The Hi-C data were aligned to the supernova assembly using *Juicer* (Durand et al. 2016). Hi-C genome assembly was performed using the *3D-DNA* pipeline (Dudchenko et al. 2017) and the output was reviewed using *Juicebox Assembly Tools* (Dudchenko et al. 2018). The Hi-C data are available on www.dnazoo.org/assemblies/Peromyscus_crinitus, where they can be visualized using *Juicebox*.*js*, a cloud-based visualization system for Hi-C data (Robinson et al. 2018).

Benchmarking Universal Single-Copy Orthologs (*BUSCO* v3, using the Mammalia odb9 database; Simão et al. 2015) and *OrthoFinder2* (Emms and Kelly 2015) were used to assess genome quality and completeness. Genome sizes were estimated for each species using *abyss-fac* (Simpson et al. 2009) and the *assemblathon_stats*.*pl* script available at: https://github.com/ucdavis-bioinformatics/assemblathon2-analysis/. *RepeatMasker* v.4.0 (Smit et al. 2015) was used to identify repetitive elements. The genome was annotated using the software package *MAKER* (3.01.02; Campbell et al. 2014). Control files, protein, and transcript data used for this process are available at https://github.com/macmanes-lab/pecr_genome/tree/master/annotation. We used *Mashmap (-f one-to-one --pi 90 -s 300000*; Jain et al. 2017, 2018) to assess syntenic conservation between *P. crinitus* and *P. maniculatus* genomes and alignments were plotted with the script *generateDotPlot*.*pl. Peromyscus crinitus* chromosomes were renamed and sorted using *seqtk* (github.com/lh3/seqtk) following the *P. maniculatus* chromosome naming scheme.

For comparative genomics analyses, we generated low-coverage whole-genome resequencing data for nine *P. crinitus* and five *P. maniculatus* individuals collected from arid sites in southern California (Fig. 1; Table S1). *Peromyscus crinitus* samples were also collected from the University of California (UC) DCDRC and *P. maniculatus* were collected further East from the UC Motte Rimrock (MOT) and Elliot Chaparral Reserves (ELL; Fig. 1). We also used publicly available low-coverage whole-genome resequencing data from 26 *P. eremicus* individuals, also collected from DCDRC and MOT, that were prepared and sequenced in parallel (Tigano et al. 2020). All samples were collected in 2009, with the exception of eight *P. eremicus* samples that were collected in 2018. Animals were collected live in Sherman traps and a 25 mg ear-clip was taken from each individual and stored at −80°C in 95% ethanol. Animals were sampled from arid areas with average monthly temperatures between 9-40°C and mean annual rainfall of 15-18 cm. The Biotechnology Resource Center at Cornell University (Ithaca, NY, USA) prepared genomic libraries using the Illumina Nextera Library Preparation kit (e.g., skim-seq). Libraries were sequenced at Novogene (Sacramento, CA, USA) using 150 bp paired-end reads from one lane on the Illumina NovaSeq S4 platform. *fastp* v. 1 (Chen et al. 2018) was used to assess read quality and trim adapters. Sequences from all samples and all species were mapped to the *P. crinitus* reference genome with *BWA* (Li and Durbin 2010) to enable comparative analyses, duplicates were removed with *samblaster v. 0*.*1*.*24* (Faust and Hall 2014), and alignments were indexed and sorted using *samtools v. 1*.*10* (Li et al. 2009).

### Population Genomics

We used the software package *ANGSD v. 0*.*93* (Korneliussen et al. 2014) to call variants from low-coverage population genomic data from the three species (26 *P. eremicus*, 9 *P. crinitus*, 5 *P. maniculatus*) with high confidence. First, an initial list of high-quality SNPs was identified by analyzing all samples from the three species together using the settings: -*SNP_pval 1e-6 - minMapQ 20 -minQ 20 -setMinDepth 20 -minInd 20 -minMaf 0*.*01*. Then, allele frequencies for each of those high-quality SNPs were calculated independently for each species, with the following filtering steps: a minimum of half (*-minInd*) *P. crinitus* and *P. eremicus* samples and all *P. maniculatus* samples had to meet independent quantity (*-minMapQ*) and quality (*-minQ*) thresholds for each variable site.

Differentiation among species was examined using a multidimensional scaling (MDS) analysis in *ANGSD*. MDS plots were generated in *R* v.3.6.1 (R Core Team 2017) based on the covariance matrix. Cook’s D was used to identify outliers (Cook and Weisberg 1984; Williams 1987). As an additional measure of differentiation, we estimated weighted and unweighted global *F*_*ST*_ values for each species pair using *realSFS* in *ANGSD. NGSadmix v. 33* (Skotte et al. 2013) was used to fit genomic data into K populations to parse species-level differences and provide a preliminary screen for genomic admixture under a maximum-likelihood model. Individuals with < 90% assignment to a particular species were considered putatively admixed. To examine the impact of coverage on the detection of admixture we also evaluated coverage distributions among admixed and non-admixed individuals. Nonetheless, expanded sample sizes with greater sequencing depth will be necessary to detail patterns of population structure and introgression. We tested K = 1 through K = (N - 1), where N is the number of total individuals examined. *NGSadmix* was run for all species combined and again for each species independently.

We used Pairwise sequential Markovian Coalescent (*PSMC v. 0*.*6*.*5-r67;* Li and Durbin 2011) to examine historical demographic changes through time for each species. *PSMC* analyses are not suitable for low-coverage genomes, therefore we used the higher-coverage reads used to generate the high-quality, chromosome-length assemblies for each species (*P. crinitus*, assembly methods detailed above; *P. eremicus*, SAMEA5799953, Tigano et al. 2020; *P. maniculatus*: GCA_003704035.1, Harvard University). Quality reads (q > 20; *Skewer*, Jiang et al. 2014) were mapped to their respective *de novo* assembled reference to identify heterozygous sites. Reference assemblies were then indexed in *BWA. Samblaster* removed PCR duplicates and *picard* (http://broadinstitute.github.io/picard/) added a read group to the resulting bam file and generated a sequence dictionary (*CreateSequenceDictionary*) from the reference assembly. For each species, *samtools* was used to sort and index alignments, and variants were called using *mpileup* in *bcftools v1*.*10*.*2* (*call*, Li et al. 2009). Consensus sequences were called in *VCFtools v 0*.*1*.*16* (*vcf2fq*, Danecek et al. 2011). *PSMC* distributions of effective population size (*N*_*e*_) were estimated with 100 bootstrap replicates and results were visualized with *gnuplot v. 5*.*2* (Williams and Kelley 2010), using perl scripts available at github.com/lh3/psmc. Output was scaled by a generation time of 6 months (0.5 yr, Millar 1989; Pergams and Lacy 2008) and a general mammalian mutation rate of 2.2 × 10^−9^ substitutions/site/year (Kumar and Subramanian 2002).

### Tests for selection & convergence

We used *Sweepfinder2* (Nielsen et al. 2005; Huber et al. 2016; DeGiorgio et al. 2016) to detect recent selective sweeps as it is compatible with low-coverage whole-genome data. This method performs a composite likelihood ratio (CLR) test to detect deviations from the neutral site frequency spectrum (SFS) that may indicate recent positive selection. *Sweepfinder2* was run on both variant and invariant sites (Huber et al. 2016) for each species, excluding sex chromosomes. Sex-chromosomes were excluded for three reasons: (1) sex chromosome evolution is both rapid and complex relative to autosomes, (2) we had different sample sizes of each sex across species, and (3) desert adaptations, the focus of this study, are unlikely to be sex-specific. We repeated *Sweepfinder2* analyses on *P. eremicus*, initially analyzed by Tigano et al. (2020), using an improved annotation scheme based on *Peromyscus-*specific data rather than *Mus musculus*. Allele frequencies were estimated in *ANGSD* independently for each species and converted to allele counts, and the site frequency spectrum (SFS) was estimated in *Sweepfinder2* from autosomes only. Identification of sweeps were based on the pre-computed SFS and the CLR was calculated every 10,000 sites. Per Tigano et al. (2020), a 10 kbp window size was selected as a trade-off between computational time and resolution. CLR values above the 99.9^th^ percentile of the empirical distribution for each species were considered to be evolving under a model of natural selection, hereafter referred to as significant sweep sites. Smaller sample sizes produce fewer bins in the SFS and a lower number of rare alleles may impact both the overall SFS and local estimate surrounding sweep sites; therefore, we explored the impact of sample sizes on *Sweepfinder2* results by downsampling the number of genomes analyzed for each species to five individuals (the total number of low-coverage genomes available for *P. maniculatus*) and compared sweep results between downsampled and all-sample datasets for the three smallest chromosomes: 21, 22, and 23.

For each species, mean Tajima’s D was calculated across the entire genome in non-overlapping windows of 10 kbp and 1 kbp in *ANGSD*. Nucleotide diversity (π) was also calculated in 10 kbp and 1 kbp windows and corrected based on the number of sites genotyped (variant and invariant) per window. Tajima’s D and π are expected to be significantly reduced in regions surrounding selective sweeps (Smith and Haigh 1974; Kim and Stephan 2002), therefore we also used a Mann-Whitney test (p < 0.05, after a Bonferroni correction for multiple tests) to measure significant deviations from the global mean in 1 kbp and 10 kbp flanking regions surrounding significant sweep sites. We also examined D and π for flanking regions surrounding 27 candidate genes identified in a previous transcriptomic investigation of *P. eremicus* and potentially involved in dehydration tolerance (MacManes 2017; Table S2). Candidate loci include aquaporins (N = 12), sodium-calcium exchangers (*SLC8a1*), and *Cyp4* genes belonging to the Cytochrome P450 gene family (N = 14). We used custom python scripts to functionally annotate (I) the closest gene to each significant sweep site, (II) the nearest upstream and downstream gene, regardless of strand (sense/antisense), and (III) the nearest upstream and downstream gene on each strand. Dataset I follows the general assumption that proximity between a significant sweep site and a protein-coding gene suggests interaction. Dataset II represents an extension of that model by encompassing the most proximal gene in each direction. Because *Sweepfinder2* is based on unphased data mapped to a consensus sequence and our data is unphased, we do not have information indicating on which strand a significant sweep site occurs. Therefore, dataset III encompasses strand-uncertainty by including the two nearest genes to a significant sweep site on both strands. It should be noted that the genes identified in smaller datasets (I, II) are nested within the larger datasets (II, III) and by definition, the larger datasets include more noise, which may dilute a signature of parallel evolution, but may better capture the true signal of selection. Hence, it is important to critically examine numerous hierarchical gene subsets. Without a linkage map, these analyses remain exploratory and can be better refined with estimates of linkage disequilibrium, linkage block sizes, and gene density in future investigations. We tested genes from each dataset for functional and pathway enrichment in Gene Ontology (GO) categories using *Panther v. 15*.*0* (Mi et al. 2017) and extracted GO terms for each enriched functional group. We used *Mus musculus* as a reference and a Bonferroni correction for multiple tests (p < 0.05) to correct for false discoveries. Enriched GO terms were summarized and visualized in *REVIGO* (Reduce and Visualize Gene Ontology, Supek et al. 2011) implemented at: http://revigo.irb.hr/index.jsp?error=expired. As a test for similar evolutionary responses to desert environments, overlap in the gene names and enriched GO terms associated with significant selective sweeps was assessed for each dataset. Overlap was visualized in the R package *VennDiagram* (Chen and Boutros 2011). To test for convergence, we used a Fisher’s Exact Test (p < 0.05) in the *GeneOverlap* package (Shen 2016) in R to assess whether gene or enriched GO term overlap between species was greater than expected based on the total number of genes/GO terms in the genome. To determine if signatures of selection were driven by differences in sequencing depth, we calculated local coverage in 10 kbp windows surrounding significant sweep sites and averaged local coverage estimates across all sweeps on a single chromosome, using a modification of: https://github.com/AdamStuckert/Ranitomeya_imitator_genome/blob/master/GenomeAssembly/DuplicateOrthologWorkbook.md. Local coverage surrounding significant sweep sites for each chromosome were compared to the chromosomal average calculated for each species (*samtools coverage --min-MQ 20, --region chr*).

To compare patterns of gene family expansion and contraction potentially involved in adaptation within the genus *Peromyscus*, we analyzed 14 additional genomes, including ten *Peromyscus* species and four near outgroup rodent species: *Microtus ochrogaster, Neotoma lepida, Sigmodon hispidus*, and *Mus musculus* (Table S3). To prevent bias driven by variable assembly qualities, samples with < 70% complete mammalian BUSCOs were excluded from downstream analyses, resulting in the final analysis of ten species (Table S3). Groups of orthologous sequences (orthogroups) were identified in *Orthofinder2*. Invariant orthogroups and groups that varied by more than 25 genes across taxa (custom python script: ortho2cafe.py) were excluded. Our rooted species tree, estimated in *Orthofinder2*, was used to calculate a single birth-death parameter (lambda) and estimate changes in gene family size using *CAFE v*.*4*.*2*.*1* (Han et al. 2013). Results were summarized using the python script *cafetutorial_report_analysis*.*py* available from the Hahn Lab at: hahnlab.github.io/CAFÉ/manual.html.

## RESULTS

### Chromosome-length genome assembly for Pcrinitus

Linked reads combined with Hi-C scaffolding produced a high-quality, chromosome-length genome assembly for *P. crinitus*. Our assembly has a contig N50 of 137,026 bp and scaffold N50 of 97,468,232 bp, with 24 chromosome-length scaffolds. The anchored sequences in the three *Peromyscus* genome assemblies were as follows: *P. crinitus* genome ∼2.27 Gb, *P. eremicus* ∼2.5 Gb, and *P. maniculatus* ∼2.39 Gb (Table 1). Our assembly has high contiguity and completeness and low redundancy, as demonstrated by the presence of 89.3% complete BUSCOs, 0.9% of duplicates, and 9.0% missing, excluding unplaced scaffolds. As anticipated based on karyotypic analyses (Smalec et al. 2019), we found no significant variation in chromosome number or major interchromosomal rearrangements between *P. crinitus* and *P. maniculatus* (Fig. S4). We annotated 17,265 total protein coding genes in the *P. crinitus* genome. Similar to other *Peromyscus* species, LINE1 (long interspersed nuclear elements) and LTR (long terminal repeats) elements comprised 22.7% of the repeats in the *P. crinitus* genome, with SINEs (short interspersed nuclear elements) representing an additional 9.6% (Table S5). Although similar to other *Peromyscus* species, *P. crinitus* has the greatest total repeat content (> 37%; see Tigano et al. 2020 Supplementary Table 2).

**Table 1.** Assembly stats, genome size, and global Tajima’s D and pi (1 kbp windows) for each *Peromyscus* species.

| Species | N | Scaffold N50 | Contig N50 | Size (Gb)<sup>a</sup> | Size (Gb)<sup>b</sup> | Taj. D | $\pi$ |
| --- | --- | --- | --- | --- | --- | --- | --- |
| *P. crinitus* | 9 | 94,816,992 | 204,461 | 2.27 | 2.28 | -0.69 | 0.005 |
| *P. eremicus* | 26 | 119,957,392 | 76,024 | 2.45 | 2.54 | -1.27 | 0.007 |
| *P. maniculatus* | 5 | 115,033,041 | 42,400 | 2.33 | 2.44 | -1.62 | 0.012 |

<sup>a</sup> abyss-fac estimate
<sup>b</sup> assemblathon estimate

### Population Genomics

MDS analysis parsed the three species into three well-separated clusters and identified no outliers or evidence of admixture (Fig. S6). *NGSadmix* identified all three species as a single group (K = 1) with the highest likelihood, but a three-population model neatly parsed the three species as expected (Fig. S7, Table S8). *NGSadmix* analysis, which is more sensitive to low sample sizes than MDS analyses, showed putative admixture in *P. crinitus* with at least three individuals displaying 11-27% ancestry from *P. eremicus* and additional material from *P. maniculatus* (4-16%). Variable samples sizes may impact assignment certainty and expanded sequencing of additional *Peromyscus* species and populations will be required to identify potential sources of introgressed material. Four *P. eremicus* individuals had < 90% assignment probability to the *P. eremicus* species cluster, with a maximum of 15% assignment to a different species cluster. Identification of admixture in both species was not biased by differences in coverage, as low (2X), medium (8X), and high coverage (17X) samples were found to be admixed at a < 90% assignment threshold (Fig. S9). No *P. maniculatus* individuals were identified as admixed.

*PSMC* estimates of historical demography (> 10 kya, Nadachowska-Brzyska et al. 2016) show greater variance and a higher overall *N*_*e*_ for *P. crinitus* relative to *P. eremicus* (Fig. 2). Demographic estimates for *P. maniculatus* are included as an additional comparison but should be interpreted with caution as they are based on sequence data from a captive-bred individual and may not accurately reflect the demography of wild populations.

**Figure 2.**
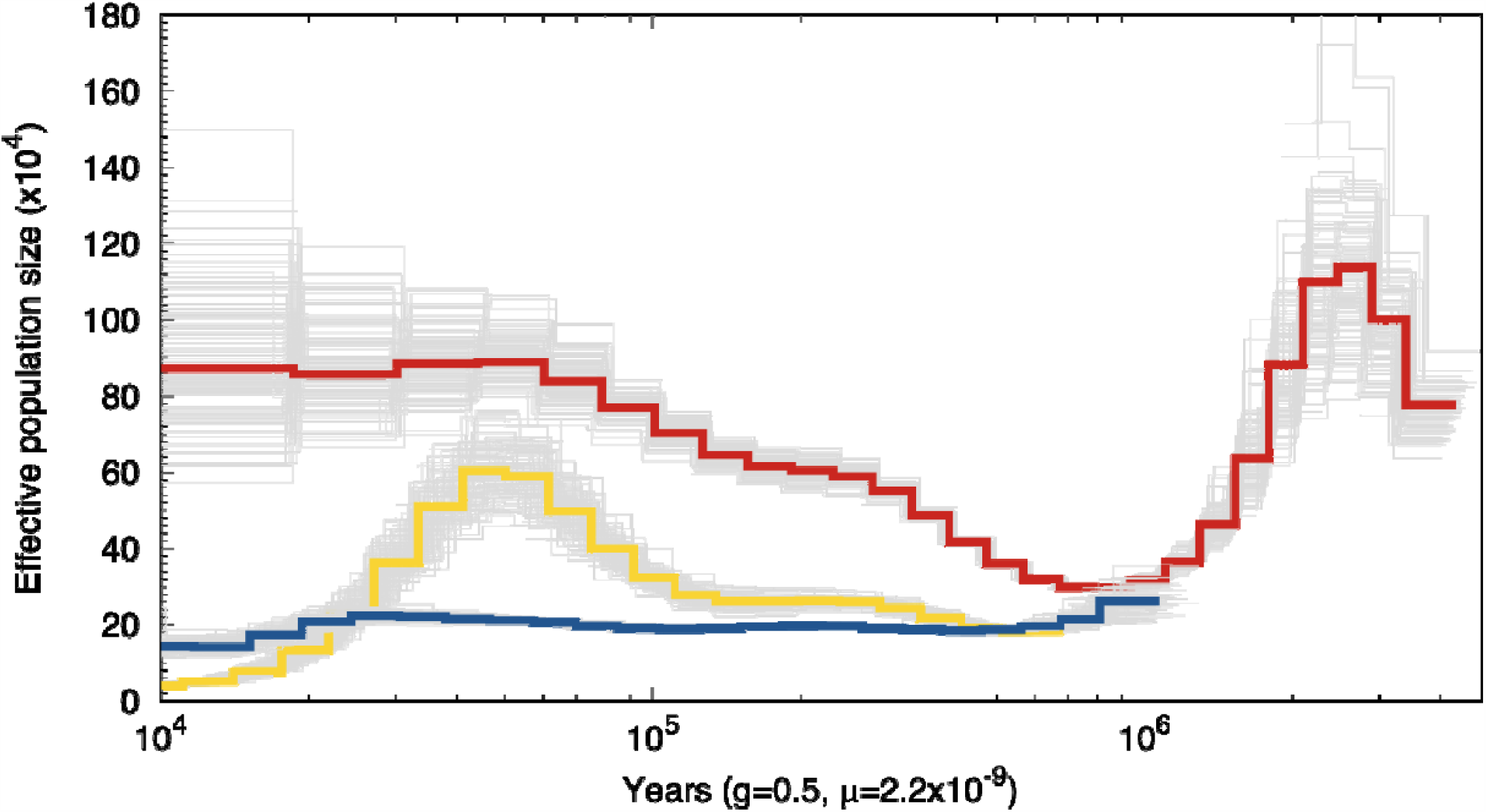
Distributions of effective population size (N_e_) through time for *P. crinitus* (red), *P. eremicus* (blue), and *P. maniculatus* (yellow) based on a generation time of 6 months (0.5 years) and a general mammalian mutation rate of 2.2×10^−9^ substitutions/site/year. Note that the *P. maniculatus* genome was sequenced from a captive individual and therefore does not reflect natural populations trends of this species.

### Selection & convergence

*Sweepfinder2* results were generally consistent across downsampled (N = 5) and all-sample datasets (9 *P. crinitus*, 26 *P. eremicus*). For example, there were no significant sweep sites identified on chromosome 21 for *P. crinitus* using either dataset, and although more sweeps were significant on chromosome 23 for the downsampled dataset (16 vs. 8), 15 of the 16 sweeps were proximal to a single protein coding gene (*S15A3*) that was also identified as experiencing a significant sweep when using all available data. All genes proximal to significant sweep sites in the downsampled *P. crinitus* dataset were also identified when all samples were analyzed. Six additional genes were identified as experiencing selective sweeps when the complete dataset was evaluated. Results for *P. eremicus* were slightly less consistent: identical numbers of sweep sites were detected on chromosomes 22 and 23 with additional significant sweep sites identified on chromosome 21 when all samples were examined. The majority of sweep sites on chromosome 21 in *P. eremicus* were proximal to *G3P*, which represented a significant sweep in both downsampled and all-sample datasets; however, there were eight additional protein coding genes proximal to significant sweep sites that were detected using the downsampled dataset, but not detected when all samples were included in the analysis (e.g., *Peripherin-2, BICRAL, Mrps18b*). We hypothesize that population structure may increase inconsistency in these results, as the 26 *P. eremicus* samples represent two populations (Motte and DCDRC) and three distinct collection events (Motte 2018, DCDRC 2009 and DCDRC 2018; Table S1), whose representation in the reduced dataset may vary due to different sample sizes and random selection of downsampled individuals. While the study design does not allow us to distinguish between type 1 and type 2 statistical errors, we hypothesize that inconsistencies are related to differences in statistical power, with greater power in the full datasets relative to the subsampled datasets.

Within *P. crinitus* we identified a total of 209 significant sweep sites (Table S10), with 104 sites localized on chromosomes 9 and 16 experiencing major selective sweeps (Fig. 3). We found 239 total significant sweep sites for *P. eremicus* (Table S11, Fig. S12). Despite the large size of chromosome 1 and strong signature of selective sweeps in *P. eremicus*, we found no significant sweep sites on this chromosome for *P. crinitus*. Finally, we identified a total of 213 significant sweep sites for *P. maniculatus* (Table S13), with 103 sites located on chromosome 4. Despite general chromosomal synteny among *Peromyscus* species (Fig. S4, Tigano et al. 2020), the chromosomal distribution of sweep sites differed among species. For example, *P. eremicus* had at least one significant sweep detected on every chromosome, while sweeps were only detected on 8 or 13 chromosomes in *P. maniculatus* and *P. crinitus*, respectively. We found a number of sweep sites were concentrated on chromosome 9 for both desert specialist species, with additional significant sweep sites for *P. crinitus* localized on chromosome 16 (Fig. 3, S12; Table S10-11). Sweeps in *P. eremicus* were widespread across the genome, with a large peak (56 significant sweep sites) on chromosome 1 (Table 11; Fig. S12). *Peromyscus maniculatus* sweeps fell primarily on chromosomes 4 and 20 (Table S13). The chromosomal distribution of significant sweep sites does not appear to be driven by differences in coverage (Table S14). Average sequencing depth for 10 kbp windows surrounding each significant sweep site did not differ significantly from the global average sequencing depth for *P. crinitus* (p = 0.25) or *P. maniculatus* (p = 0.28). Local (10 kbp window) coverage surrounding significant sweeps for *P. eremicus* were less consistent (Table S10, S11, S13; Supplementary Results). Fourteen of 232 significant sweep sites (6%) in *P. eremicus* exhibited extreme local sequencing depths of 0 or >1,000, leading to a significant difference between overall mean sequencing depth and sequencing depth surrounding sweep sites (p = 0.03), with sweep windows exhibiting higher sequencing depths on average (73 vs. 25). If the 14 anomalous values are excluded, sequencing coverage does not differ (p = 0.37) for *P. eremicus*.

**Figure 3.**
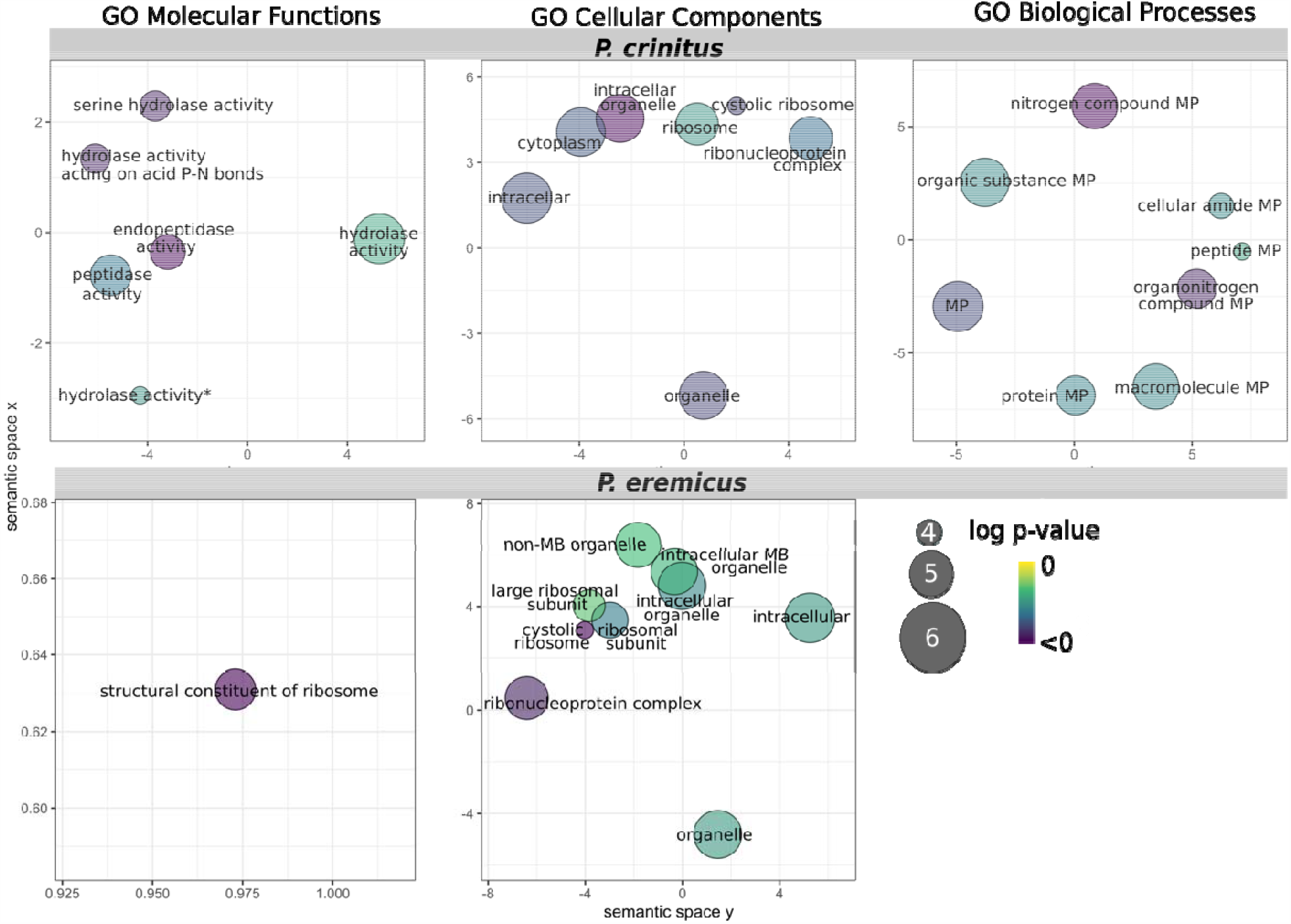
*REVIGO* plots of enriched functional groups for *P. crinitus* (top row) and *P. eremicus* (bottom row) based on functional annotation of the two nearest protein-coding genes to each site (dataset II) identified as the subject of a selective sweep. Darker colors indicate greater significance. MP = metabolic process, MB = membrane-bound.

The effect of a selective sweep extends beyond the specific site identified as the target of positive selection; hence, putative outliers (CLR > 99.9%) are indicative of a sweep in the 10 kbp window but the specific nucleotides under selection cannot be identified. As coding genes represent only a small proportion of the genome, if a sweep site does not fall in one of these regions we assume that the target of selection may be a regulatory element affecting gene expression of proximal coding genes. Under this assumption, we hierarchically examined protein coding genes most proximal to each sweep site and chose not to set a distance threshold, as regulatory elements are known to affect genes up to hundreds of kb away (e.g., Wallbank et al. 2016). On average the distance from a sweep site to the nearest coding gene was 45 kbp in *P. crinitus* (range: 31 - 439,122 bp, median = 5 kbp) and much greater for both *P. maniculatus* (average: 152 kbp; range: 190 - 513,562 bp, median = 111 kbp) and *P. eremicus* (average: 117 kbp; range: 38 - 1,423,027 bp; median: 35 kbp), despite high assembly qualities for all species and identical methods of gene annotation. For both *P. eremicus* and *P. maniculatus*, only two significant sweep sites were localized within protein-coding genes (*P. eremicus:* Meiosis-specific with OB domain-containing protein, Harmonin; *P. maniculatus:* Dehydrogenase/reductase SDR family member 7B and Zinc finger protein 217; Table 2). In contrast, for *P. crinitus* 12 significant sweep sites fell within 19 distinct candidate loci, many of which code for multiple alternatively spliced transcripts (Table 2). Among the significant sweep sites localized within *P. crinitus* coding sequences, we identified 19 enriched GO terms (3 Biological Process [BP], 9 Molecular Function [MF], 7 Cellular Component [CC]), with functionality ranging from ‘proteolysis’ to ‘hydrolase’ activity (Fig. 4; Table S15). Functional examination of candidate loci identified solute regulation as a key function, with genes pertaining to calcium (*Trypsin-2* [*PRSS2*]) and zinc (*Kallikrein-4* [*KLK4*]) binding and sodium regulation (*Prostasin* [*PRSS8*]) indicated as under selection.

**Table 2.**
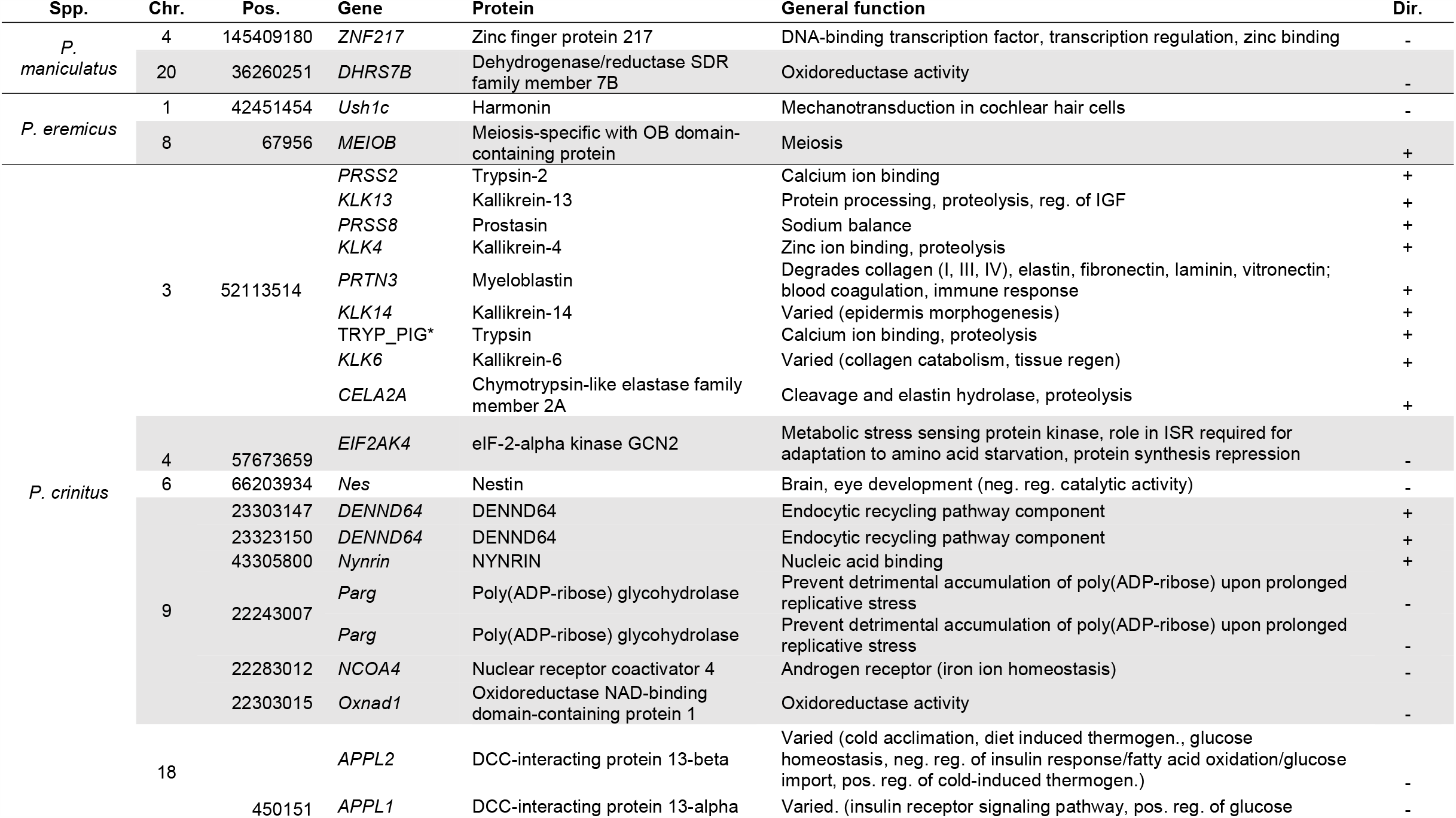
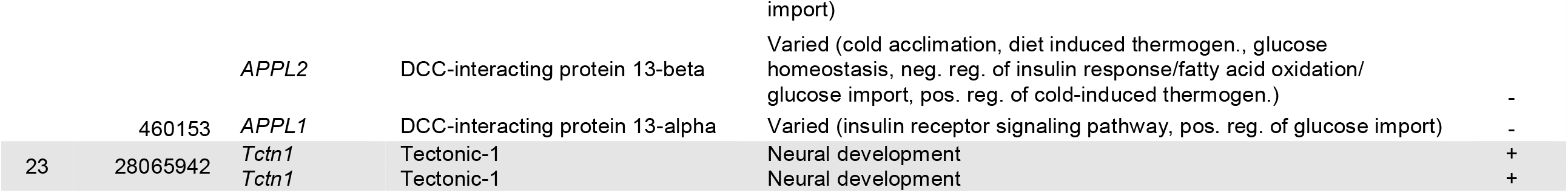
Significant sweep sites localized within protein-coding genes for each *Peromyscus* species. Species (Spp.), chromosome (Chr.) sweep position (Pos.), gene name, protein, general function (based on UniProt database: uniprot.org), and direction (Dir.) of gene transcription. Abbreviation definitions: thermogen./thermogenesis, neg./negative, pos./positive, reg./regulation, IGF/insulin-like growth factor, ISR/Integrated Stress Response. *no gene name alternative available.

**Figure 4.**
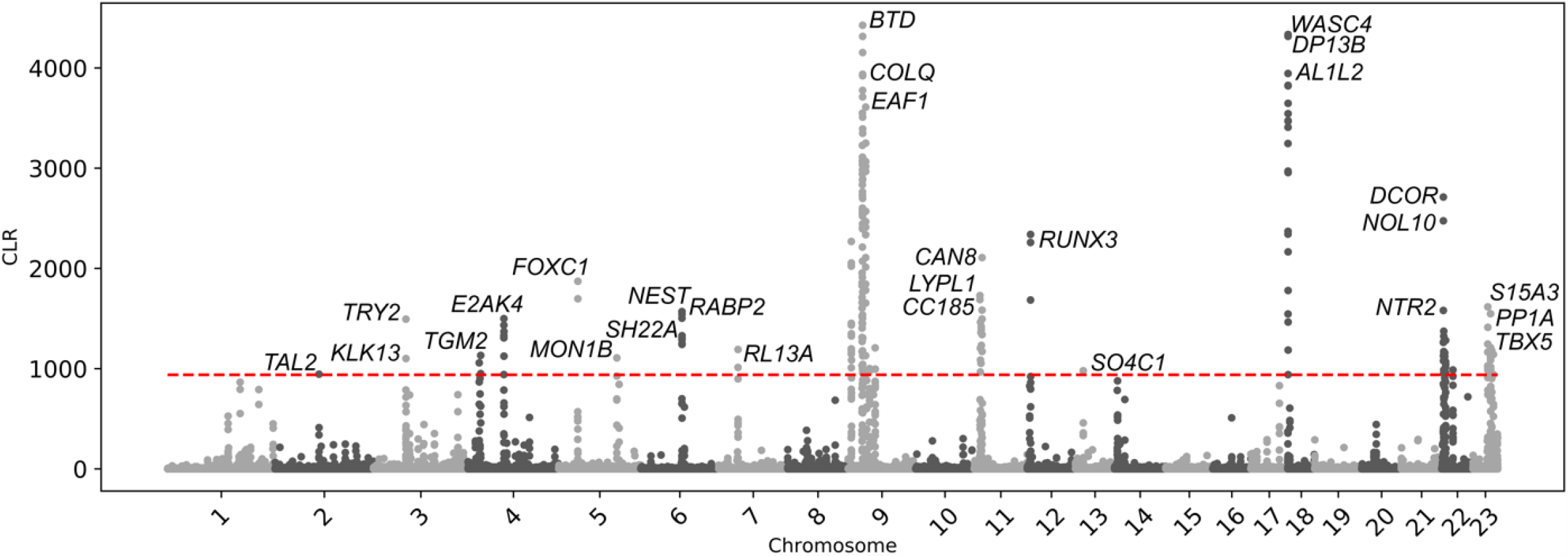
Composite likelihood ratio (CLR) scores for *P. crinitus* based on *Sweepfinder2* results. Values above the horizontal red line surpass the 99.9^th^ percentile. The top five or fewer unique genes are labeled for each chromosome.

Here we report the results for dataset II, as this dataset (i) ensures the inclusion of the most proximal gene under selection, by including the most proximal gene on each strand, and (ii) reduces noise associated with dataset III, which includes four genes proximal to each sweep site. Results for datasets I and III are addressed in more detail in the Supplementary Information. Examination of dataset II in *P. crinitus* identified 121 unique genes and 26 enriched GO terms (8 Biological Processes [BP], 10 Molecular Functions [MF], 8 Cellular Component [CC]), with functionality pertaining to metabolism (e.g., ‘protein metabolic process’, ‘organonitrogen compound metabolic process’, ‘peptide metabolic process’) and ribosomes (Fig. 4; Tables S10, S15). For *P. eremicus*, we identified 202 unique genes and 14 enriched GO terms (0 BP, 1 MF, 13 CC) associated with selective sweeps, with functionality centered around ribosomes (Table S11, S16). For *P. maniculatus*, we identified 215 unique genes and eight enriched GO terms (0 BP, 1 MF, 7 CC) associated with selective sweeps (Table S13, S17). Two genes and seven enriched GO terms that were proximal to sweep sites were shared between the two desert specialists, but the number of shared genes was not significantly different from what is expected by chance alone. Functional enrichment of *P. eremicus* and *P. maniculatus* across all datasets was limited to ribosomes (e.g., ‘structural constituent of ribosome’, ‘cytosolic ribosome’, ‘ribosomal subunit’; Fig. 4; Table 3, S15-17). In contrast, functionality of enriched GO terms for *P. crinitus* centered on metabolic processes, including protein breakdown, hydrolysis, and cellular functionality (e.g., ‘organelle’, ‘intracellular’, ‘cytoplasm’; Fig. 4; Table S15), in addition to ribosomes.

**Table 3.**
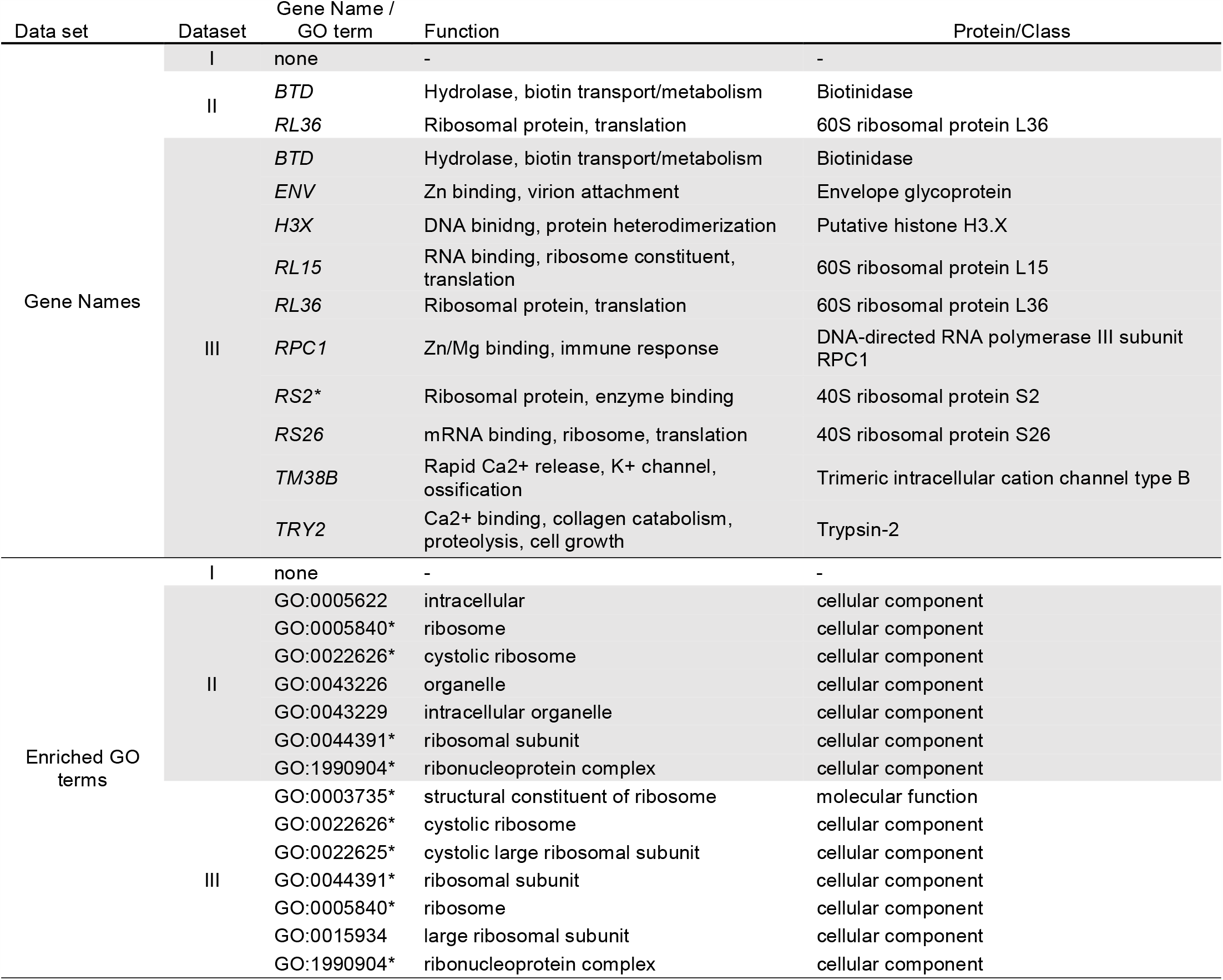
Functional annotation of proximal gene names and enriched GO (gene ontology) terms associated with significant selective sweeps and shared between desert-adapted *P. crinitus* and *P. eremicus*. * indicates gene names or GO terms also shared with *P. maniculatus*

*Peromyscus eremicus* and *P. maniculatus* shared significant overlap (p < 0.05) in enriched GO terms across all hierarchical data subsets (I, II, III; Fig. 5). Significant overlap of enriched GO terms was also detected between *P. crinitus* and both other *Peromyscus* species for datasets II and III only, with no overlap detected for dataset I (Fig. 5). Significant overlap between desert specialists *P. eremicus* and *P. crinitus* was only detected in dataset III. Overall, GO terms and genes associated with ribosomal functionality were frequently shared among all species examined, but a unique pattern of selection was not shared among desert specialists.

**Figure 5.**
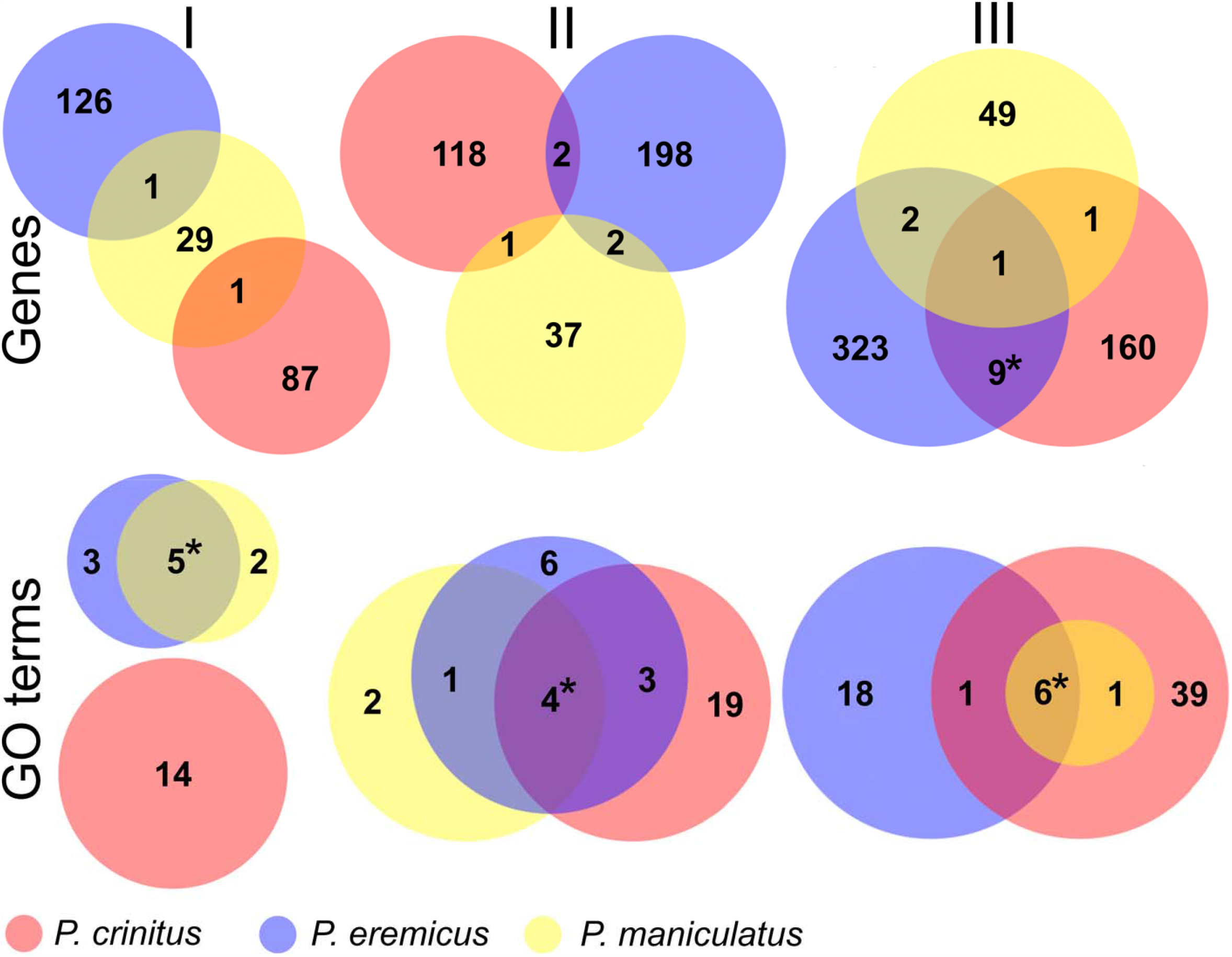
Overlap in proximal gene names (top row) and enriched GO terms (bottom row) for datasets I (left column), II (center), and III (right). *indicates significant overlap between species

Species tree estimates (Fig. S18) were consistent with previous phylogenetic investigations (Bradley et al. 2007). *Peromyscus crinitus* and *P. eremicus* are sister in our species tree, but note that a number of intermediate taxa remain unsampled (e.g., *P. merriami, P. californicus*). Among the species examined here, the two desert specialists are part of a larger clade of desert adapted *Peromyscus* to the exclusion of *P. maniculatus*. The *maniculatus* clade is comprised of *P. leucopus, P. polionotus*, and *P. maniculatus*, and the *nasutus-attwateri* clade is most basal within *Peromyscus* (Fig. S18), consistent with Platt et al. (2015). For the *Peromyscus* genus, we found 19,925 gene families that had experienced contractions, 502 expansions, and 12 families that were evolving rapidly. However, we found no gene families experiencing significant expansions, contractions, or rapid evolution below the genus level.

Average Tajima’s D (1 kbp windows) was negative for all species and ranged from −0.69 to −1.61. *Peromyscus crinitus* had the lowest Tajima’s D value and *P. maniculatus* the highest (Fig. S19-20). Global pairwise *F*_*ST*_ between species ranged from 0.20-0.27 (unweighted: 0.12-0.17). Mean global π (1 kbp windows) was 0.005 (±0.005) for *P. crinitus*, 0.007 (± 0.007) for *P. eremicus*, and 0.012 (± 0.010) for *P. maniculatus* (Fig. S21). Both Tajima’s D and π for 1 and 10 kbp flanking regions surrounding significant selective sweep sites were significantly higher than the global average for each species (Table S20). Only in *P. maniculatus* did we detect a significant reduction in π surrounding significant sweep sites. Tajima’s D for flanking regions surrounding the *a priori* candidate loci identified by MacManes (2017) were also significantly more positive in all three species (Table S20).

## DISCUSSION

Continued and accelerating environmental change increases the exigency of accurately predicting species responses to anthropogenic climate change. Adaptive evolutionary responses vary among species and populations, even when subjected to similar environmental selective pressures (Bi et al. 2015; Garcia-Elfring et al. 2019). Evidence of parallel *de novo* molecular changes or selective retention of shared ancestral variation can highlight genes or genomics regions, including but not limited to functional variants, haplotypes, or structural features of the genome, that may be key to adaptation. Alternatively, the same adaptive phenotype can evolve through alternative evolutionary strategies. Thus, it is possible, even among related species, that adaptation to similar environmental conditions will not exhibit similar patterns of molecular evolution despite similar adaptive phenotypes. We analyzed genome-wide patterns of selective sweeps among three species of deer mice within the North American genus *Peromyscus* to identify candidate loci involved in heat- and dehydration-tolerance. We hypothesized that the desert specialists, *P. crinitus* and *P. eremicus*, would share genes or pathways associated with selective sweeps that were not shared with phylogenetically ancestral *P. maniculatus*. These patterns would be indicative of parallel selection (and therefore, parallel evolution) on either *de novo* mutations or ancestral variation. Given the suite of desert adaptations shared by these species, shared signatures of selection may relate to survival in high-temperature, low-water environments. Additionally, we hypothesized that shared patterns of selective sweeps and enriched functional groups across all three species, if present, would highlight candidate loci underpinning local adaptation of *P. maniculatus* to arid conditions and potentially identify common loci involved in the repeated evolution of desert adaptation. Although the species examined here are monophyletic, the two desert specialists share a more recent ancestor (Fig. S18) and there are number of unsampled taxa that phylogenetically separate the desert specialists from *P. maniculatus* (e.g., *P. merriami, P. californicus*). For this reason, we cannot distinguish between parallel and convergent evolution, and without evidence of ancestral divergence followed by reconvergence, we will discuss shared signatures of selection as parallel evolution hereafter.

Overall, we did not find support for parallel evolution among desert specialist species, but we identified a number of candidate loci that may be important to desert adaptation in *P. crinitus*. Instead of a shared mechanism of heat- and dehydration tolerance, we hypothesize that the two desert specialists examined here may have adapted to similar environments through divergent molecular mechanisms, with *P. crinitus* potentially responding through genomic changes to protein coding genes and *P. eremicus* through transcriptional regulation of gene expression. This hypothesis is based on the lack of overlap in selective sweeps between desert specialists, the proximity of sweeps to protein coding genes in *P. crinitus* relative *to P. eremicus*, and previous gene expression results for *P. eremicus* (Kordonowy and MacManes 2017; MacManes 2017). Molecular flexibility of thermoregulatory responses may have catalyzed the radiation of *Peromyscus* in North America by enabling rapid exploitation of novel thermal environments. Finally, the application of an evolutionary lens to the interpretation of genomic patterns of selection, particularly one that integrates historical demography and gene flow, can help parse varied evolutionary mechanisms (parallel vs. convergent, genomic vs. transcriptomic) of molecular adaptation.

### Limited evidence of parallel evolution

Identification of similar genes or functional groups under selection in different species adapted to similar environments can provide evidence in support of parallel evolution. In contrast, we found limited evidence of parallel evolution among desert-adapted *Peromyscus*. Few to no enriched GO terms overlapped between desert specialists (Fig. 5). Only GO terms relating to ribosomes (e.g., ‘ribosome’, ‘ribosomal subunit’, ‘cystolic ribosome’, etc.) overlapped between all three *Peromyscus* species examined, with the most significant overlap in GO terms occurring between *P. eremicus* and *P. maniculatus*. Although *P. maniculatus* are not generally xerocoles, the individuals sequenced here were collected in arid regions of southern California (subspecies *P. m. sonoriensis*). Therefore, the shared signature of selection on ribosomes across all examined *Peromyscus*, whether it reflects parallel evolution or the selective retention of shared ancestral polymorphisms, may be associated with adaptation to hot and dry conditions or more broadly relate to thermoregulatory plasticity among *Peromyscus* rodents. Few genes proximal to selective sweeps were shared among all species, with only one instance of significant overlap: ten genes were shared between the two desert specialists under dataset III (Fig. 5). Although dataset III may be confounded by excess noise through the inclusion of additional protein coding genes, this signature is potentially consistent with a parallel evolution. Again, many of the genes shared between *P. crinitus* and *P. eremicus* are directly related to ribosomal functionality (e.g., *RL36, RS26, RL15*) and also shared with *P. maniculatus*. Determining whether these sweeps are the result of shared new mutations or ancestral variation and whether selection on ribosomal functionality is unique to desert-adapted taxa or more broadly relevant to the genus will require additional tests for selection and expanded taxonomic sampling across the genus *Peromyscus*.

Cellular damage accumulates quickly in desert environments as a consequence of increased thermal- and osmotic-stress (Lamitina et al. 2006; Burg et al. 2007). In response, expression changes modulate osmoregulation by removing and replacing damaged proteins to prevent cell death (Lamitina et al. 2006); hence, ribosomes, which play a critical role in protein synthesis and degradation, are central to thermoregulatory responses (Porcelli et al. 2015). Although we did not find significantly expanded or contracted gene families within the genus *Peromyscus*, previous investigations of the entire Myodonta clade within Rodentia identified multiple expanded or contracted gene families associated with ribosomes in *P. eremicus* (Tigano et al. 2020). Here, ribosomes appear to be a potential target of parallel evolution in desert-adapted *Peromyscus*, yet this genomic signature is not unique to this genus, nor to desert-adapted species. First, the relative abundance of ribosome-associated genes throughout the genome (>1000 GO annotations pertaining to ribosomes, Bult et al. 2019) may intrinsically increase the representation of this functional group, especially at coarse resolution (10 kbp windows). Second, selection on ribosomal functionality may be commonly experienced across many species adapted to distinct thermal environments (metazoans; Porcelli et al. 2015). Ribosomes are evolutionarily linked to the mitochondrial genomes of animals (Barreto and Burton 2012; Bar-Yaacov et al. 2012) and accelerated mitochondrial evolution in animals has led to compensatory, rapid evolution of ribosomal proteins (Osada and Akashi 2012; Barreto and Burton 2013; Bar-Yaacov et al. 2012). Rapid mitochondrial diversification within *Peromyscus* (Riddle et al. 2000; Bradley et al. 2007; Platt et al. 2015), coincident with the ecological radiation of this genus (Lindsey 2020), suggests that equivalent, recent selection on ribosomal proteins may be a key evolutionary innovation that enabled Peromyscine rodents to successfully and quickly adapt to varied thermal environments. Alternatively, broad selection on ribosomes across all species may also contribute to other, varied aspects of these species biology. Comparisons among additional *Peromyscus* species will be necessary to test these hypotheses in detail.

Evaporative cooling through sweating, panting, or salivating increases water loss and challenges osmoregulatory homeostasis in a hot and dry climate (McKinley et al. 2018). Thermal stress exacerbates dehydration by increasing evaporative water loss and if untreated, can lead to cognitive dysfunction, motor impairment, and eventually death. In consequence, osmoregulatory mechanisms are often under selection in extreme thermal environments (MacManes and Eisen 2014; Marra et al. 2014). Consistent with the importance of osmoregulation in desert species, four of the ten protein-coding genes that experienced a significant selective sweep and were shared between desert specialist species (dataset III) are involved in ion balance (Table 3). Proteins Trypsin-2 (*TRY2*) and Trimeric intracellular cation channel type-B (*TM38B*) are associated with sweeps in both desert specialists and are involved in calcium ion (Ca^2+^) binding and release, respectively. DNA-directed RNA polymerase III (*RPC1*) has also experienced a significant sweep in both desert specialists and influences magnesium (Mg^2+^) binding. Calcium and magnesium cations are among those essential for osmoregulation (also, Na^+^, K^+^, Cl^-^, HCO_3_^-^; Stockham and Scott 2008) and parallel selection on these genes is consistent with the hypothesis that solute-carrier proteins are essential to maintaining homeostasis in desert-specialized rodents (Marra et al. 2014; Kordonowy and MacManes 2017). Additional genes involved in osmoregulation were identified as experiencing selective sweeps only in *P. crinitus* (Table 2; Table S10). Prostatin (*PRSS8*), only found to be under selection in *P. crinitus*, is critically responsible for increasing the activity of epithelial sodium (Na^+^) channels, which mediate sodium reabsorption through the kidneys (Narikiyo et al. 2002). Two more genes associated with Ca^2+^ regulation ([*PRSS2* and *TRYP*) and other genes regulating zinc (*KLK4*) and iron (*NCOA4*) were also identified as targets of selective sweeps exclusively in *P. crinitus*.

Genomic scans for selective sweeps based on the SFS are only one way to detect signatures of parallel evolution and these methods can be sensitive to missing data, including low-coverage and small sample sizes; thus, the putative roles of these candidate genes in desert adaptation remains to be explored using additional methods with increased sequencing depth (see Booker et al. 2017, Weigand and Leese 2018) and other experimental approaches (e.g., MacManes 2017).

### Metabolic tuning: proteins-for-water or lipids-for-torpor?

*f* contrast to the protein-for-water Hot deserts experience dramatic fluctuations in both food and water availability that challenge species survival (Noy-Meir 1973; Silanikove 1994). Mammals accommodate high temperatures by increasing body temperatures, to a point, and cold temperatures by aerobic thermogenesis or metabolic suppression via the initiation of torpor or hibernation (Levesque et al. 2016). When resources are scarce, metabolism relies exclusively on endogenous nutrients; carbohydrates (e.g., sugars, glucose) are consumed immediately, then lipids, and eventually, proteins. Protein oxidation has a low-energy return relative to lipid catabolism (Bar and Volkoff 2012), but yields five times more metabolic water (Jenni and Jenni-Eiermann 1998; Gerson and Guglielmo 2011a, b; McCue et al. 2017). Therefore, in a low-water environment an early shift to protein catabolism during periods of resource limitation may represent an important water source for desert species (e.g., protein-for-water hypothesis; Mosin 1984; Jenni and Jenni-Eiermann, 1998; Gerson and Guglielmo, 2011a, b). Consistent with this hypothesis, we identified numerous candidate genes that experienced selective sweeps in *P. crinitus* and that are involved in the detection of metabolic-stress and shifts in metabolic fuel consumption. For example, the gene eIF-2-alpha kinase GCN2 (*E2AK4*), which is responsible for sensing metabolic stress in mammals and required for adaptation to amino acid starvation, experienced the strongest selective sweep on chromosome 4 in *P. crinitus* (Fig. 3; Harding et al. 2003; Baker et al. 2012; Taniuchi et al. 2016). Numerous candidate genes involved in oxidation (Oxidoreductase NAD-binding domain-containing protein 1 [*Oxnad1*]), fat catabolism (Kallikrein-6 [*KLK6*]), protein processing (Kallikrein-13 [*KLK13*]), and proteolysis (Kallikrein [*KLK4, KLK13*], Trypsin [*PRSS2, TRYP, TRY2*], Chymotrypsin-like elastase family member 2A [*CELA2A*]) were associated with significantly enriched GO terms in *P. crinitus*. Proteolysis was the most enriched functional group in *P. crinitus* (Fig. 4; Table S15), potentially supporting the protein-for-water hypothesis.

In contrast to the protein-for-water hypothesis, efficient lipid acquisition and storage may be critical to enabling heat- and drought-induced torpor (Buck et al. 2002; Melvin and Andrews 2009), which allows long duration, low energy survival in desert adapted species, including *Peromyscus*. Significant weight loss in experimentally-dehydrated *P. eremicus* and enhanced thermogenic performance of high-altitude-adapted deer mice have been associated with enhanced lipid metabolism (Cheviron et al. 2012; Kordonowy et al. 2016). At high altitudes, increased lipid oxidation enables aerobic thermogenesis, but in hot deserts lipids may represent a valuable energy source in a food-scarce environment (e.g., lipids-for-torpor hypothesis). Two additional candidate genes, DCC-interacting protein 13-alpha and -beta (*APPL1, APPL2*), experienced significant selective sweeps in *P. crinitus* and are important in glucose regulation, insulin response, and fatty acid oxidation, potentially supporting the lipids-for-torpor hypothesis. Laboratory manipulations of *APPL1* demonstrate protection against high-fat diet-induced cardiomyopathy in rodents (Park et al. 2013) and *APPL2* is responsible for dietary regulation, cold-induced thermogenesis, and cold acclimation (uniprot.org). Together, these genes play a role in both obesity and dietary regulation. Both *APPL* genes are associated with obesity and non-alcoholic fatty liver disease and their sweep signature in *P. crinitus* has relevant connections to biomedical research that remain to be explored (Jiang et al. 2012; Barbieri et al. 2013). Physiological tests will be essential to determine whether desert-adapted deer mice prioritize proteins or fats during periods of resource limitation (e.g., lipids-for-torpor) or extreme dehydration (e.g., protein-for-water hypothesis).

Molecular rewiring of metabolic processes in response to environmental conditions has been documented in a number of species (e.g., mammals, Velotta et al. 2020; birds, Xie et al. 2018; fruit flies, Mallard et al. 2018), but expression changes can also impact species metabolism (Cheviron et al. 2012; Storz and Cheviron 2016). The capacity for rapid molecular adaptation to distinct thermal environments through either transcriptomic regulation or changes to protein coding genes, combined with thermoregulatory behavioral fine-tuning (e.g., nocturnality, aestivation, food caching, burrowing, dietary shifts), suggests there may be many evolutionary strategies available for small mammals to accommodate increasing temperatures. Anthropogenic change, however, is occurring at a rate that far outpaces the evolutionary timescales on which these adaptations have naturally evolves; Thus, while metabolism and metabolic plasticity represent fundamental phenotypes for anticipating species survival under altered climate scenarios natural selection, alone they may be insufficient for species survival.

### Different evolutionary strategies, same result

Diverse functional enrichment of the *P. crinitus* genome (Fig. 4), spanning metabolic and osmoregulatory functions, in addition to the general functional enrichment of ribosomes, identifies a number of candidate loci worthy of detailed examination. Additional, comparisons across populations and environments will illustrate the influence of these loci and others in thermoregulation, dehydration, and other adaptive traits. Significant selective sweeps that are not shared among desert specialists, including most of the loci detected here, may still be related to desert adaptation but could also be related to other aspects of this species biology.

There are multiple evolutionary routes to achieve environmental adaptation, most notably through genomic changes in protein coding genes or transcriptional regulation of gene expression. Lack of evidence for parallel evolution between desert specialists, the proximity of significant selective sweeps to protein coding genes, diverse functional enrichment of *P. crinitus* relative to *P. eremicus*, and previous gene expression results for *P. eremicus* (Kordonowy and MacManes 2017; MacManes 2017) lead us to hypothesize alternative evolutionary strategies for each desert specialist, each shaped by their independent demographic histories: *P. crinitus* primarily through changes in protein coding genes and *P. eremicus* primarily through transcriptional regulation. Evidence of many significant sweep sites in the *P. eremicus* genome, located more distant from protein coding genes, and with functional enrichment restricted to ribosomes, suggests that adaptation in this species may be driven more by selection on regulatory or non-coding regions of the genome that impact gene expression, a hypothesis that is consistent with transcriptomic investigations in this species (MacManes and Eisen 2014; Kordonowy and MacManes 2017) and other *Peromyscus* and rodents (Cheviron et al. 2012; Marra et al. 2014; Storz and Cheviron 2016). Without equivalent gene expression data for *P. crinitus*, we cannot eliminate a similarly important role for transcriptional regulation and look forward to testing this hypothesis in greater detail with RNAseq data. Transcriptional regulation is a particularly useful mechanism for environmental acclimation, as these changes are more transient relative to genomic changes and can enhance phenotypic flexibility (Garrett and Rosenthal 2012; Rieder et al. 2015; Liscovitch-Brauer et al. 2017). Reduced variation is expected near selective sweeps and can encompass tens to thousands of adjacent nucleotides depending on recombination and the strength of selection (Fay and Wu 2000; Carlson et al. 2005), yet counter to expectations, Tajima’s D and nucleotide diversity for regions flanking putative selective sweeps were significantly higher than the global average for most comparisons (Table S20). The same observation, elevated Tajima’s D and nucleotide diversity surrounding selective sweeps, was also made in *P. eremicus* (Tigano et al. 2020). This counterintuitive pattern holds across different window sizes (1 kbp, 10 kbp) and warrants further investigation.

Placing the results of selective sweep analyses within an evolutionary framework is also critical to interpreting adaptive evolutionary responses. The deer mouse genus *Peromyscus* originated approximately 8 Mya, followed by a massive radiation around 5-6 Mya that led to the divergence of a monophyletic clade now comprised of desert adapted taxa, although the ancestral state of this clade remains unknown. These desert adapted species may have colonized arid environments through either a single or multiple invasions, with further interspecific divergence thereafter (Platt et al. 2015). The expansion of North American deserts following the conclusion of the last glacial maximum (∼11 Kya; Pavlik et al. 2008) constrains the adaptive timescales of contemporary desert species. The consistently stable and low historical effective population size of *P. eremicus* suggests that his species has harbored less genetic variation for selection to act on, despite similar levels of contemporary diversity relative to *P. crinitus* (Fig. 2). In consequence, adaptive evolution of *P. eremicus* is likely to have been more impacted by genetic drift (Allendorf 1986; Masel 2011) relative to *P. crinitus*, which historically has a larger effective population size and therefore a broader pool of variation for selection to act upon across evolutionary timescales, which could explain the higher diversity of genes and enriched GO terms compared to *P. eremicus*. Within this context, changes in regulatory elements that mediate gene expression may have been a more efficient means of environmental adaptation available to *P. eremicus* (Allendorf 1986; Neme and Tautz 2016; Mallard et al. 2018), whereas the larger historical effective population size of *P. crinitus* is more conducive to the maintenance of higher levels of genetic diversity and may have enabled the rapid evolution of protein coding sequences through mutational stochasticity, the reduced impact of genetic drift, a larger pool of standing genetic variation, and potentially, gene flow. *Peromyscus crinitus* experienced a historical demographic bottleneck prior to the formation of North American deserts; Nevertheless, the recovered effective population size of *P. crinitus* is much larger than *P. eremicus* and consistent with low levels of detected admixture in *P. crinitus* (Fig. S7, Table S8). Negative Tajima’s D values can indicate population expansion following a bottleneck, consistent with both the demographic history of *P. crinitus* and putative admixture in this species. Evidence of a historical bottleneck is also reinforced by moderate to high levels of nucleotide diversity in *P. crinitus*. Repeated growth and contraction of rivers in the American Southwest during Pleistocene glacial-interglacial cycles (0.7-0.01 Mya; Muhs et al. 2003; Van Dam and Matzke 2016) would have provided iterative opportunities for connectivity and introgression between incompletely-isolated *Peromyscus* species. Historical hybridization between *P. crinitus* and one or more other Peromyscine species, likely unsampled here, may have accelerated adaptation in *P. crinitus* through the rapid influx of novel mutation combinations through adaptive introgression, a hypothesis that warrants further investigation through expanded taxonomic sampling and explicit tests of adaptive introgression. Low-coverage whole-genome resequencing is optimal for population genomics investigations (O’Rawe et al. 2015; da Fonseca et al. 2016), but limits detailed analyses of historical introgression. We look forward to testing this hypothesis with expanded population sampling and increased sequencing depth. Finally, linkage disequilibrium decay is also weaker in larger populations, where recombination is higher, therefore it’s possible that the shorter distance between significant sweep sites and the nearest coding gene in *P. crinitus* could be due to the larger historical population sizes of this species relative to *P. eremicus*. However, the evolutionary scales of *PSMC* and *Sweepfinder2* do not overlap, *PSMC* characterizes historical demography beyond 10 kya, whereas selective sweeps have occurred recently. Overall, incorporating an evolutionary perspective into the interpretation of selection patterns has important implications for understanding species responses to changing climate, as historical demography and gene flow, in addition to selection, shape genetic diversity over evolutionary timescales.

## Conclusion

Contrasting patterns of selective sweeps and evolutionary histories between different species experiencing similar environmental pressures can provide powerful insights into the adaptive potential of species. We used comparative and population genomic analyses of three *Peromyscus* species to identify candidate loci that may underlie adaptations to desert environments. Candidate loci identified in *P. crinitus* serve to inform future investigations focused on predicting potential for adaptation and identifying the causes of warming-related population declines (Cahill et al. 2013). The identification of numerous targets of selection within *P. crinitus* highlights multiple molecular mechanisms (metabolic switching, osmoregulatory tuning) associated with physiological responses to deserts that warrant further investigation.. Our approach demonstrates the importance of placing genomic selection analyses into an evolutionary framework to anticipate evolutionary responses to change.

## ACKNOWLEDGEMENTS

We thank A. S. Westbrook for computational support; the Premise computing cluster at the University of New Hampshire, where all analyses were conducted; the Biotechnology Resource Center at Cornell University for preparation of whole-genome resequencing libraries; Christopher Tracy for access to the Boyd Deep Canyon Reserve; Jim Patton for desert field expertise; Sen Pathak, Asha Multani, Richard Behringer for providing the fibroblast samples from the T.C. Hsu Cryo-Zoo at the University of Texas M.D. Anderson Cancer Center; DNA Zoo for generating Hi-C data; Pawsey Supercomputing Centre with funding from the Australian Government and the Government of Western Australia for computational support of the DNA Zoo effort and the Museum of Southwestern Biology at the University of New Mexico for loaned tissue materials; two anonymous reviewers whose comments significantly improved the manuscript. This work was funded by the National Institute of Health National Institute of General Medical Sciences to MDM (1R35GM128843).

## DATA AVAILABILITY STATEMENT

The draft assembly data are housed on the European Nucleotide Archive (ENA) under project ID PRJEB33592. The Hi-C data is available on SRA (<u>SRX7041777, SRX7041776</u>, <u>SRX7041773</u>) under the DNA Zoo project accession <u>PRJNA512907</u>. The *P. crinitus* genome assembly is available at https://www.dnazoo.org/assemblies/Peromyscus_crinitus. Whole-genome resequencing data for *P. crinitus* are available on ENA under project ID PRJEB35488. Custom python scripts and other bash scripts used in analysis are available at: https://github.com/jpcolella/Peromyscus_crinitus.

## REFERENCES

Abbott, K. (1971). Water economy of the canyon mouse, Peromyscus crinitus stephensi. Comparative Biochemistry and Physiology, 38A: 37–52.

Allendorf, F. (1986). Genetic drift and the loss of alleles versus heterozygosity. Zoo Biology, 5: 181–190.

Anderson, K., & Jetz, W. (2005). The broad-scale ecology of energy expenditure of endotherms. Ecology Letters, 8: 310–318.

Baker, B., Nargund, A., Sun, T., & Haynes, C. (2012). Protective coupling of mitochondrial function and protein synthesis via the eIF2α kinase GCN-2. PLoS Genetics, 8: e1002760.

Bar, N., & Volkoff, H. (2012). Adaptation of the physiological, endocrine and digestive system functions to prolonged food deprivation in fish. In Comparative physiology of fasting, starvation, and food limitation (pp. 69–90). New York: Springer.

Bar-Yaacov, D., Blumberg, A., & Mishmar, D. (2012). Mitochondrial-nuclear co-evolution and its effects on OXPHOS activity and regulation. Biochimica et Biophsyica Acta (BBA)-Gene Regulatory Mechanisms, 1819: 1107–1111.

Barbieri, M., Esposito, A., Angellotti, E., Rizzo, M., Marfella, R., & Paolisso, G. (2013). Association of genetic variation in adaptor protein APPL1/APPL2 loci with non-alcoholic fatty liver disease. PLOS ONE, 8: e71391.

Barreto, F., & Burton, R. (2013). Evidence for compensatory evolution of ribosomal proteins in response to rapid divergence of mitochondrial rRNA. Molecular Biology and Evolution, 30: 310–314.

Bassham, S., Catchen, J., Lescak, E., von Hippel, F., & Cresko, W. (2018). Repeated selection of alternatively adapted haplotypes creates sweeping genomic remodeling in stickleback. Genetics, 209: 921–939.

Bedford, M., & Hoekstra, H. (2015). The natural history of model organisms: Peromyscus mice as a model for studying natural variation. Elife, 4: e06813.

Bi, K., Linderoth, T., Singhal, S., Vanderpool, D., Patton, J., Nielsen, R., … Good, J. (2015). Temporal genomic contrasts reveal rapid evolutionary responses in an alpine mammal during recent climate change. PLoS Genetics, 15: e1008119.

Booker, T., Jackson, B., & Keightley, P. (2017). Detecting positive selection in the genome. BMC Biology, 15. doi:https://doi.org/10.1186/s12915-017-0434-y

Bradley, R., Durish, N., Rogers, D., Miller, J., Engstrom, M., & Kilpatrick, C. (2007). Toward a molecular phylogeny for Peromyscus: Evidence from mitochondrial cytochrome-b sequences. Journal of Mammalogy, 88: 1146–1159.

Buck, M., Squire, T., & Andrews, M. (2002). Coordinate expression of PDK4 gene: a means of regulating fuel selection in a hibernating mammal. Physiological Genomics, 8: 5–13.

Bult, C. J., Blake, J. A., Smith, C. L., Kadin, J. A., Richardson, J. E., and the Mouse Genome Database Group. (2019). Mouse Genome Database (MGD). Nucleic Acids Research, 8: D801–D806.

Burg, M., Ferraris, J., & Dmitrieva, N. (2007). Cellular response to hyperosmotic stresses. Physiological Reviews, 87: 1441–1474.

Cahill, A., Aiello-Lammens, M., Fisher-Reid, M., Hua, X., Karanewsky, C., Yeong Ryu, H., … Wiens, J. (2013). How does climate change cause extinction? Proceedings of the Royal Society B: Biological Sciences, 280: 20121890.

Campbell, M., Holt, C., Moore, B., & Yandell, M. (2014). Genome annotation and curation using MAKER and MAKER-P. Current Protocols in Bioinformatics, 48: 4–11.

Carlson, C., Thomas, D., Eberle, M., Swanson, J., Livingston, R., Rieder, M., & Nickerson, D. (2005). Genomic regions exhibiting positive selection identified from dense genotype data. Genome Research, 15: 1553–1565.

Charlesworth, B. (2009). Effective population size and patterns of molecular evolution and variation. Nature Reviews Genetics, 10: 195–205.

Chen, H., & Boutros, P. (2011). VennDiagram: a package for the generation of highly-customizable Venn and Euler diagrams in R. BMC Bioinformatics, 12: 35

Chen, I., Hill, J., Ohlemueller, R., Roy, D., & Thomas, C. (2011). Rapid range shifts of species associated with high levels of climate warming. Science, 333: 1024–1026.

Chen, S., Zhou, Y., Chen, Y., & Gu, H. (2018). fastp: an ultra-fast all-in-one FASTQ preprocessor. Bioinformatics, 34: i884–i890.

Cheviron, Z., Bachman, G., Connaty, A., McClelland, G., & Storz, J. (2012). Regulatory changes contribute to the adaptive enhancement of thermogenic capacity in high-altitude deer mice. Proceedings of the National Academy of Sciences, 109: 8635–8640.

Cheviron, Z., Connaty, A., McClelland, G., & Storz, J. (2014). Functional genomics of adaptation to hypoxic cold-stress in high-altitude deer mice: transcriptomic plasticity and thermogenic performance. Evolution, 68: 48–62.

Cook, R., & Weisberg, S. (1984). Residuals and influence in Regression. Wiley.

Coyne, J. A., & Orr, H. A. (2004). Speciation. Sunderland, MA: Sinauer Associates.

da Fonseca, R. R., Albrechtsen, A., Themudo, G., Ramos-Madrigal, J., Sibbesen, J., Maretty, L., … Pereira, R. (2016). Next-generation biology: sequencing and data analysis approaches for non-model organisms. Marine Genomics, 30: 3–13. doi:10.1016/j.margen.2016.04.012

Danecek, P., Auton, A., Abecasis, G., Albers, C. A., Banks, E., DePristo, M. A., … 1000 Genome Project Data Process Subgroup. (2011). The variant call format and VCFtools. Bioinformatics, 27: 2156–2158.

Degen, A. (2012). Ecophysiology of small desert mammals. Springer Science & Business Media.

DeGiorgio, M., Huber, C., Hubisz, M., Hellmann, I., & Nielsen, R. (2016). SweepFinder2: increased sensitivity, robustness and flexibility. Bioinformatics, 32: 1895–1897.

Dewey, M., & Dawson, W. (2001). Deer mice: “the Drosophila of North American mammalogy. Genesis, 29: 105–109.

Dudchenko, O., Batra, S., Omer, A., Nyquist, S., Hoeger, M., Durand, N., … Aiden, E. (2017). De novo assembly of the Aedes aegypti genome using Hi-C yields chromosome-length scaffolds. Science, 356: 92–95.

Dudchenko, O., Shamim, M., Batra, S., Durand, N., Musial, N., Mostofa, R., … Aiden, E. (2018). The Juicebox Assembly Tools module facilitates de novo assembly of mammalian genomes with chromosome-length scaffolds for under $1000. BioRxiv. doi:https://doi.org/10.1101/254797

Durand, N., Shamim, M., Machol, I., Rao, S., Huntley, M., Lander, E., & Aiden, E. (2016). Juicer provides a one-click system for analyzing loop-resolution Hi-C experiments. Cell Systematics, 3: 95–98.

Emms, D. M., & Kelly, S. (2015). OrthoFinder: solving fundamental biases in whole genome comparisons dramatically improves orthogroup inference accuracy. Genome Biology, 16: 157.

Faust, G., & Hall, I. (2014). SAMBLASTER: fast duplicate marking and structural variant read extraction. Bioinformatics, 30: 2503–2505.

Fay, J., & Wu, C.-I. (2000). Hitchhiking under positive Darwinian selection. Genetics, 155: 1405–1413.

Freeman, B., Lee-Yaw, J., Sunday, J., & Hargreaves, A. (2018). Expanding, shifting and shrinking: The impact of global warming on species’ elevational distributions. Global Ecology and Biogeography, 27: 1268–1276.

Fumagalli, M., Vieira, F., Linderoth, T., & Nielsen, R. (2014). ngsTools: methods for population genetics analyses from next-generation sequencing data. Bioinformatics, 30: 1486–1487.

Garcia-Elfring, A., Barrett, R., & Millien, V. (2019). Genomic signatures of selection along a climatic gradient in the northern range margin of the white-footed mouse (Peromyscus leucopus). Journal of Heredity, 110: 684–695.

Garrett, S., & Rosenthal, J. (2012). RNA editing underlies temperature adaptation in K+ channels from polar octopuses. Science, 335: 848–851.

Gerson, A., & Guglielmo, C. (2011a). Flight at low ambient humidity increases protein catabolism in migratory birds. Science, 333: 1434–1436.

Gerson, A., & Guglielmo, C. (2011b). House sparrows (Passer domesticus) increase protein catabolism in response to water restriction. American Journal of Physiology, 300: R925– R930.

Glazier, D. (1980). Ecological shifts and the evolution of geographically restricted species of North American Peromyscus (mice). Journal of Biogeography, 7: 63–83.

Han, M., Thomas, G., Lugo-Martinez, J., & Hahn, M. (2013). Estimating gene gain and loss rates in the presence of error in genome assembly and annotation using CAFE 3. Molecular Biology and Evolution, 30: 1987–1997.

Harding, H., Zhang, Y., Zeng, H., Novoa, I., Lu, P., Calfon, M., … Ron, D. (2003). An integrated stress response regulates amino acid metabolism and resistance to oxidative stress. Molecular Cell, 11: 619–633.

Hoffmann, A., & Sgrò, C. (2011). Climate change and evolutionary adaptation. Nature, 470: 479–485.

Hoffmann, A., & Willi, Y. (2008). Detecting genetic responses to environmental change. Nature Reviews: Genetics, 9: 421–432.

Hu, C., & Hoekstra, H. (2017). Peromyscus burrowing: A model system for behavioral evolution. Seminars in Cell and Developmental Biology, 61: 107–114.

Huber, C., DeGiorgio, M., Hellmann, I., & Nielsen, R. (2016). Detecting recent selective sweeps while controlling for mutation rate and background selection. Molecular Ecology, 25: 142– 156.

Issaian, T., Urity, V., Dantzler, W., & Pannabecker, T. (2012). Architecture of vasa recta in the renal inner medulla of the desert rodent Dipodomys merriami: potential impact on the urine concentrating mechanism. American Journal of Physiology - Regulatory, Integrative and Comparative Physiology, 303: R748–R756.

Ivy, C., & Scott, G. (2017). Control of breathing and ventilatory acclimatization to hypoxia in deer mice native to high altitudes. Acta Physiologica, 221: 266–282.

Jain, C., Koren, S., Dilthey, A., Phillippy, A. M., Aluru, S. (2018) A fast adaptive algorithm for computing whole-genome homology maps. Bioinformatics, 34: i748–i756.

Jain, C., Dilthey A, Koren, S, Aluru, S., Phillippy, A. M. (2017) A fast approximate algorithm for mapping long reads to large reference databases. In: Sahinalp S. (eds) Research in Computational Molecular Biology. RECOMB 2017. Lecture Notes in Computer Science, 10229. Springer, Cham.

Jenni, L., & Jenni-Eiermann, S. (1998). Fuel supply and metabolic constraints in migrating birds. Journal of Avian Biology, 29: 521–528.

Jiang, H., Lei, R., Ding., S.-W., Zhu, S. (2014). Skewer: a fast and accurate adapter trimmer for next-generation sequencing paired-end reads. BMC Bioinformatics, 15: 182.

Jiang, S., Fang, Q., Yu, W., Zhang, R., Hu, C., Dong, K., … Jia, W. (2012). Genetic variations in APPL2 are associated with overweight and obesity in a Chinese population with normal glucose tolerance. BMC Medical Genetics, 13: 22.

Johnson, D., & Armstrong, D. (1987). Peromyscus crinitus. Mammalian Species, 287: 1–8.

Jones, M., Mills, L., Alves, P., Callahan, C., Alves, J., Lafferty, D., … Good, J. (2018). Adaptive introgression underlies polymorphic seasonal camouflage in snowshoe hares. Science, 360: 1355–1358.

Kaseloo, P., Crowell, M., & Heideman, P. (2014). Heritable variation in reaction norms of metabolism and activity across temperatures in a wild-derived population of white-footed mice (Peromyscus leucopus). Journal of Comparative Physiology B, 184(4): 525–534.

Kim, Y., & Stephan, W. (2002). Detecting a local signature of genetic hitchhiking along a recombining chromosome. Genetics, 160: 765–777.

Kordonowy, L., & MacManes, M. (2017). Characterizing the reproductive transcriptomic correlates of acute dehydration in males in the desert-adapted rodent, Peromyscus eremicus. BMC Genomics, 18: 473.

Kordonowy, L., Lombardo, K., Green, H., Dawson, M., Bolton, E., LaCourse, S., & MacManes, M. (2016). Physiological and biochemical changes associated with acute experimental dehydration in the desert adapted mouse, Peromyscus eremicus. Physiological Reports, 5: e13218.

Korneliussen, T., Albrechtsen, A., & Nielsen, R. (2014). ANGSD: Analysis of Next Generation Sequencing Data. Bioinformatics, 15: 356.

Kumar, S., & Subramanian, S. (2002). Mutation rates in mammalian genomes. Proceedings of the National Academy of Sciences, 99: 803–808.

Lamitina, T., Haung, C., & Strange, K. (2006). Genome-wide RNAi screening identifies protein damage as a regulator of osmoprotective gene expression. Proceedings of the National Academy of Sciences of the United States of America, 103: 12173–12178.

Levesque, D., Nowack, J., & Stawski, C. (2016). Modelling mammalian energetics: the heterothermy problem. Climate Change Responses, 3: 7.

Li, H., & Durbin, R. (2010). Fast and accurate long-read alignment with Burrows-Wheeler transform. Bioinformatics, 26: 589–595.

Li, Heng, & Durbin, R. (2011). Inference of human population history from whole genome sequence of a single individual. Nature, 475: 493–496.

Li, H., Handsaker, B., Wysoker, A., Fennel, T., Ruan, J., Homer, N., … 1000 Genome Project Data Process Subgroup. (2009). The sequence alignment/map format and SAMtools. Bioinformatics, 25: 2078–2079.

Lindsey, L. (2020). Utilizing genomic applications to examine patterns of diversification in deermice (Rodentia: Cricetidae: Peromyscus). Texas Tech Dissertation.

Liscovitch-Brauer, N., Alon, S., Porath, H., Elstein, B., Unger, R., Ziv, T., … Eisenberg, E. (2017). Trade-off between transcriptome plasticity and genome evolution in Cephalopods. Cell, 169: 191–202.

MacManes, M., & Eisen, M. (2014). Characterization of the transcriptome, nucleotide sequence polymorphism, and natural selection in the desert adapted mouse Peromyscus eremicus. PeerJ, 2: e642.

MacManes, M. (2017). Severe acute dehydration in a desert rodent elicits a transcriptional response that effectively prevents kidney injury. American Journal of Physiology. Renal Physiology, 313: F262–F272.

MacMillen, R., & Christopher, E. (1975). The water relations of two populations of non-captive desert rodents. In Environmental physiology of desert organisms ((N. F. Hadley, ed.): pp. 117–137). Stroudsburg, Pennsylvania: Dowden, Hutchinson, and Ross.

MacMillen, R. (1972). Water economy of nocturnal desert rodents. Symposia of the Zoological Society of London, 31: 147–174.

MacMillen, R. (1983). Water regulation in Peromyscus. Journal of Mammalogy, 64: 38–47.

Mallard, F., Nolte, V., Tobler, R., Kapun, M., & Schlötterer, C. (2018). A simple genetic basis of adaptation to a novel thermal environment results in complex metabolic rewiring in Drosophila. Genome Biology, 19: 1–15. doi:10.1186/s13059-018-1503-4

Marra, N., Romero, A., & DeWoody, J. (2014). Natural selection and the genetic basis of osmoregulation in heteromyid rodents as revealed by RNA-seq. Molecular Ecology, 23: 2699–2711.

Masel, J. (2011). Genetic drift. Current Biology, 21: R837–R838.

McCue, M., Sandoval, J., Beltran, J., & Gerson, A. (2017). Dehydration causes increased reliance on protein oxidation in mice: a test of the protein-for-water hypothesis in a mammal. Physiological and Biochemical Zoology, 90: 359–369.

McDonald, M., Gehrig, S., Meintjes, P., Zhang, X.-X., & Rainey, P. (2009). Adaptive divergence in experimental populations of Pseudomonas fluorescens. IV. Genetic constraints guide evolutionary trajectories in a parallel adaptive radiation. Genetics, 183: 1041–1053.

McKinley, M., Martelli, D., Pennington, G., Trevaks, D., & McAllen, R. (2018). Integrating competing demands of osmoregulatory and thermoregulatory homeostasis. Physiology, 33: 170–181.

McNab, B. (1963). The influence of fat deposits on the basal rate of metabolism in desert homoiotherms. Comparative Biochemistry and Physiology, 26: 337–343.

McNab, B., & Morrison, P. (1963). Body temperature and metabolism in subspecies of Peromyscus from arid and mesic environments. Ecological Monographs, 33: 63–82.

Melvin, R., & Andrews, M. (2009). Torpor induction in mammals: Recent discoveries fueling new ideas. Trends in Endocrinology and Metabolism, 20: 490–498.

Mi, H., Huang, X., Muruganujan, A., Tang, H., Mills, C., Kang, D., & Thomas, P. (2017). PANTHER version 11: expanded annotation data from Gene Ontology and Reactome pathways, and data analysis tool enhancements. Nucleic Acids Research, 45: F183–D189.

Millar, J. (1989). Reproduction and development. In Advances in the Study of Peromyscus (Rodentia) (eds Kirkland Gl Jr, Layne JN, pp. 169–232). Lubbock, Texas: University of Texas Press.

Morhardt, J., & Hudson, J. (1966). Daily torpor induced in white-footed mice (Peromyscus spp.) by starvation. Nature, 212: 1046–1047.

Moritz, C., Patton, J., Conroy, C., Parra, J., White, G., & Beissinger, S. (2008). Impact of a century of climate change on small-mammal communities in Yosemite National Park, USA. Science, 322: 261–264.

Morjan, C., & Rieseberg, L. (2004). How species evolve collectively: implications of gene flow and selection for the spread of advantageous alleles. Molecular Ecology, 13: 1341–1356.

Mosin, A. (1984). On the energy fuel in voles during their starvation. Comparative Biochemistry and Physiology, 77: 563–565.

Muhs, D., Reynolds, R., Been, J., & Skipp, G. (2003). Eolian sand transport pathways in the southwestern United States: importance of the Colorado river and local sources. Quaternary International, 104: 3–18.

Murray, G. G. R., Soares, A. E. R., Novak, B. J., Schaefer, N. K., Cahill, J. A., Baker, A. J., … Shapiro, B. (2017) Natural selection shaped the rise and fall of passenger pigeon genomic diversity. Science 17: 951–954.

Nadachowska-Brzyska, K., Burri, R., Linnéa, S., Hans, E. (2016). PSMC analysis of effective population sizes in molecular ecology and its application to black-and-white Ficedula flycatchers. Molecular Ecology 25: 1058–1072.

Narikiyo, T., Kitamura, K., Adachi, M., Miyoshi, T., Iwashita, K., Shiraishi, N., … Tomita, K. (2002). Regulation of prostasin by aldosterone in the kidney. The Journal of Clinical Investigation, 109: 401–408.

Natarajan, C., Hoffmann, F., Lanier, H., Wolf, C., Cheviron, Z., Spangler, M., … Storz, J. (2015). Intraspecific polymorphism, interspecific divergence, and the origins of function-altering mutations in deer mouse hemoglobin. Molecular Biology and Evolution, 32: 978–997.

Neme, R., & Tautz, D. (2016). Fast turnover of genome transcription across evolutionary time exposes entire non-coding DNA to de novo gene emergence. ELife, 5: e09977.

Nielsen, R., Williamson, S., Kim, Y., Hubisz, M., Clark, A., & Bustamante, C. (2005). Genomic scans for selective sweeps using SNP data. Genome Research, 15: 1566–1575.

Noy-Meir, I. (1973). Desert ecosystems: Environment and producers. Annual Review of Ecology and Systematics, 4: 25–51.

O’Rawe, J., Ferson, S., & Lyon, G. (2015). Accounting for uncertainty in DNA sequencing data. Trends in Genetics, 31: 61–66.

Osada, N., & Akashi, H. (2012). Mitochondrial-nuclear interactions accelerated compensatory evolution: evidence from the primate cytochrome C oxidase complex. Molecular Biology and Evolution, 29: 337.

Park, M., Wu, D., Park, T., Choi, C., Li, R.-K., Cheng, K., … Sweeney, G. (2013). APPL1 transgenic mice are protected from high-fat diet-induced cardiac dysfunction. American Journal of Physiology: Endocrinology and Metabolism, 305: E795–E804.

Pavlik, B. M. (2008). The California deserts: An ecological rediscovery. Berkeley: University of California Press.

Pergams, O., & Lacy, R. (2008). Rapid morphological and genetic change in Chicago-area Peromyscus. Molecular Ecology, 17: 450–463.

Pierce, S., & Vogt, F. (1993). Winter acclimatization in Peromyscus maniculatus gracilis P leucopus noveboracensis, and P. l. leucopus. Journal of Mammalogy, 74: 665–677.

Platt, II, R., Amman, B., Keith, M., Thompson, C., & Bradley, R. (2015). What is Peromyscus? Evidence from nuclear and mitochondrial DNA sequences suggests the need for a new classification. Journal of Mammalogy, 96: 708–719.

Porcelli, D., Butlin, R., Gaston, K., Joly, D., & Snook, R. (2015). The environmental genomics of metazoan thermal adaptation. Heredity, 114: 502–514.

R Core Team. (2017). R: A language and environment for statistical computing. Vienna, Austria: R Foundation for Statistical Computing.

Riddle, B., Hafner, D., & Alexander, L. (2000). Phylogeography and systematics of the Peromyscus eremicus species group and the historical biogeography of North American warm regional deserts. Molecular Phylogenetics and Evolution, 17: 145–160.

Rieder, L., Savva, Y., Reyna, M., Chang, Y.-J., Dorsky, J., Rezaei, A., & Reenan, R. (2015). Dynamic response of RNA editing to temperature in Drosophila. BMC Biology, 13: 1.

Robinson, J., Turner, D., Durand, N., Thorvaldsdottir, H., Mesirov, J., & Aiden, E. (2018). Juicebox.js provides a cloud-based visualization system for Hi-C data. Cell Systems, 6: 256– 258.

Rosenblum, E., Parent, C., & Brandt, E. (2014). The molecular basis of phenotypic convergence. Annual Review of Ecology and Systematics, 45: 203–226.

Rundle, H., Nagel, L., Boughman, J., & Schluter, D. (2000). Natural selection and parallel speciation in sympatric sticklebacks. Science, 287: 306–308.

Schwimmer, H., & Haim, A. (2009). Physiological adaptations of small mammals to desert ecosystems. Integrative Zoology, 4: 357–366.

Shen, L. (2016). GeneOverlap: R package for testing and visualizing gene overlaps. New York City, New York: Ichan School of Medicine at Mount Sinai.

Sikes, R. S., & The Animal Care and Use Committee of the American Society of Mammalogists. (2016). 2016 Guidelines of the American Society of Mammalogists for the use of wild mammals in research and education. Journal of Mammalogy, 97: 663–688.

Silanikove, N. (1994). The struggle to maintain hydration and osmoregulation in animals experiencing severe dehydration and rapid rehydration: the story of ruminants. Experimental Physiology, 79: 281–300.

Simão, F. A., Waterhouse, R. M., Ioannidis, P., Kriventseva, E. V., & Zdobnov, E. M. (2015). BUSCO: assessing genome assembly and annotation completeness with single-copy orthologs. Bioinformatics, 31: 32-10–3212.

Simpson, J., Wong, K., Jackman, S., Schein, J., Jones, S., & Birol, I. (2009). ABySS: a parallel assembler for short read sequence data. Genome Research, 19: 1117–1123.

Skotte, L., Korneliussen, T., & Albrechtsen, A. (2013). Estimating individual admixture proportions from next generation sequencing data. Genetics, 195: 693–702.

Smalec, B., Heider, T., Flynn, B., & O’Neill, R. (2019). A centromere satellite concomitant with extensive karyotypic diversity across the Peromyscus genus defies predictions of molecular drive. Chromosome Research: An International Journal on the Molecular, Supramolecular and Evolutionary Aspects of Chromosome Biology. doi:10.1007/s10577-019-09605-1

Smit, A., Hubley, R., & Green, P. (2013). RepeatMasker Open-4.0. Available at: http://www.repeatmasker.org/

Smith, J., & Haigh, J. (1974). The hitch-hiking effect of a favourable gene. Genetical Research, 23: 23–35.

Steppan, S., Adkins, R., & Anderson, J. (2004). Phylogeny and divergence-date estimates of rapid radiations in Muroid rodents based on multiple nuclear genes. Systematic Biology, 53: 533–553.

Stockham, S., & Scott, M. (2008). Fundamental of veterinary clinical pathology. Ames, IA: Wiley-Blackwell.

Storz, J., & Cheviron, Z. (2016). Functional genomic insights into regulatory mechanisms of high-altitude adaptation. Advances in Experimental Medicine and Biology, 903: 113–128. doi:10.1007/978-1-4988-7678-9_8.

Storz, J., & Kelly, J. (2008). Effects of spatially varying selection on nucleotide diversity and linkage disequilibrium: insights from deer mouse globin genes. Genetics, 180: 367–379.

Storz, J. (2007). Hemoglobin function and physiological adaptation to hypoxia in high-altitude mammals. Journal of Mammalogy, 88: 24–31.

Storz, J., Runck, A., Moriyama, H., Weber, R., & Fago, A. (2010). Genetic differences in hemoglobin function between highland and lowland deer mice. Journal of Experimental Biology, 213: 2565–2574.

Storz, J., Cheviron, Z., McClelland, G., & Scott, G. (2019). Evolution of physiological performance capacities and environmental adaptation: insights from high-elevation deer mice (Peromyscus maniculatus). Journal of Mammalogy, 100: 910–922.

Supek, F., Bošnjak, M.,Škunca, N., & Šmuc, T. (2011). REVIGO summarizes and visualizes long lists of gene ontology terms. PloS One, 6: e21800.

Taniuchi, S., Miyake, M., Tsugawa, K., Oyadormari, M., & Oyadomari, S. (2016). Integrated stress response of vertebrates is regulated by four eIF2α kinases. Scientific Reports, 6: 32886.

Tigano, A., Colella, J., & MacManes, M. (2020). Comparative and population genomics approaches reveal the basis of adaptation to deserts in a small rodent. Molecular Ecology, 29: 1300–1314.

Tigano, A. & Friesen, V. L. (2016). Genomics of local adaptation with gene flow. Molecular Ecology, 25: 2144–2164.

Tingley, M., & Beissinger, S. (2013). Cryptic loss of montane avian richness and high community turnover over 100 years. Ecology, 94: 598–609.

Urban, M. (2015). Accelerating extinction risk from climate change. Science, 348: 571–573.

Van Dam, M., & Matzke, N. (2016). Evaluating the influence of connectivity and distance on biogeographical patterns in the south-western deserts of North America. Journal of Biogeography, 43: 1514–1532.

Veal, R., & Caire, W. (1979). Peromyscus eremicus. Mammalian Species, 118: 1–6.

Velotta, J., Robertson, C., Schweizer, R., McClelland, G., & Cheviron, Z. (2020). Adaptive shifts in gene regulation underlie a developmental delay in thermogenesis in high-altitude deer mice. Molecular Biology and Evolution, msaa086. doi:https://doi.org/10.1093/molbev/msaa086

Wagner, G., & Lynch, V. (2008). The gene regulatory logic of transcription factor evolution. Trends in Ecology & Evolution, 23: 277–385.

Wallbank, R. W., Baxter, S. W., Pardo-Diaz, C., Hanly, J. J., Martin, S. H., Mallet, J.,Dasmahapatra, K. K., Salazar, C., Joron, M., Nadeau, N., McMillan, W. O. (2016). Evolutionary novelty in a butterfly wing pattern through enhancer shuffling. PLoS biology, 14(1): p.e.1002353.

Weigand, H., & Leese, F. (2018). Detecting signatures of positive selection in non-model species using genomic data. Zoological Journal of the Linnean Society, 184: 528–583.

Weisenfeld, N. I., Kumar, V., Shah, P., Church, D. M., & Jaffe, D. B. (2017). Direct determination of diploid genome sequences. Genome Research, 27: 757–767.

Wichman, H., & Lynch, C. (1991). Genetic variation for seasonal adaptation in Peromyscus leucopus: nonreciprocal breakdown in a population cross. Journal of Heredity, 82: 197–204.

Williams, T., & Kelley, C. (2010). Gnuplot 4.4: an interactive plotting program (Version 4.4). Retrieved from http://gnuplot.sourceforge.net/

Williams, D. (1987). Generalized linear model diagnostics using the deviance and single casedeletions. Applied Statistics, 36: 181–191.

Xie, S., Yang, X., Wang, D., Zhu, F., Yang, N., Hou, Z., & Ning, Z. (2018). Thyroid transcriptome analysis reveals different adaptive responses to cold environmental conditions between two chicken breeds. PLOS ONE, 13: 2018.

